# A chromatin modulator sustains self-renewal and enables differentiation of postnatal neural stem and progenitor cells

**DOI:** 10.1101/554535

**Authors:** Kushani Shah, Gwendalyn D. King, Hao Jiang

## Abstract

It remains unknown whether H3K4 methylation, an epigenetic modification associated with gene activation, regulates fate determination of the postnatal neural stem and progenitor cells (NSCs and NPCs, or NSPCs). Here we show that the Dpy30 subunit of the major H3K4 methyltransferase complexes is preferentially expressed at a high level in NSCs and NPCs. By genetically inactivating Dpy30 in specific regions of mouse brain, we demonstrate a crucial role of efficient H3K4 methylation in maintaining both the self-renewal and differentiation capacity of postnatal NSPCs. Dpy30 inactivation results in deficiency in global H3K4 methylation, and disrupts development of hippocampus and especially the dentate gyrus and subventricular zone, the major regions for postnatal NSC activities. By in vitro assays on neurospheres from mouse brains as well as human and mouse NPCs, we show that Dpy30 is indispensable for sustaining the self-renewal and proliferation of NSPCs in a cell-intrinsic manner, and also enables the differentiation of mouse and human NPCs to neuronal and glial lineages. Dpy30 directly regulates H3K4 methylation and the induction of several genes critical in neurogenesis. These findings link a prominent epigenetic mechanism of gene expression to the fundamental properties of NSPCs, and may have implications in neurodevelopmental disorders.

**SIGNIFICANCE STATEMENT:** As a highly prominent epigenetic mark that is associated with gene activation and a number of neurodevelopmental disorders in human, the role of histone H3K4 methylation in the fate determination of neural stem cells is unclear. Result of this study uncover a profound role of this epigenetic modification in the fundamental properties of postnatal neural stem cells, including self-renewal and differentiation, and may have implications for a better understanding and treatment of a broad spectrum of neurodevelopmental disorders associated with H3K4 methylation modulators.

## INTRODUCTION

The capacity of self-renewal and differentiation is the defining property of all stem cells including neural stem cells (NSCs) [1, 2]. The maintenance and alteration of stem cell fate are ultimately controlled at the level of gene expression, which is profoundly shaped by the global and local chromatin state. Epigenetic mechanisms, including histone methylation, are increasingly linked to NSC fate determination [3-5]. Histone H3K4 and H3K27 methylation marks are a paradigm of two oppositely acting epigenetic mechanisms on gene expression [6, 7]. Both the writer and eraser of H3K27 methylation regulates the self-renewal and/or differentiation of NSCs [8-10]. Bmi1, a Polycomb family protein involved in linking H3K27 methylation to gene repression, is required for NSC self-renewal partly through repressing *Ink4a/Arf* [11, 12]. On the other hand, the role of H3K4 methylation in NSC fate determination is relatively less understood, despite being a prominent mark generally associated with and functionally important for gene activation [13-16]. Mutations of several writers, erases, and readers of this chromatin modification are associated with some human neurodevelopmental disorders [17], but a role of this modification in these disease processes is unknown. In mammalian cells, H3K4 methylation is mainly catalyzed by the Set1/Mll complexes, which comprise one of the Setd1a, Setd1b, Kmt2a/Mll1, Kmt2b/Mll2, Kmt2c/Mll3, and Kmt2d/Mll4 proteins as the catalytic subunit and all of the Wdr5, Rbbp5, Ash2l, and Dpy30 proteins as the common core subunits [18, 19]. Among these subunits, Mll1 has been shown to be required for postnatal neurogenesis, but not gliogenesis [20]. However, no reduction in H3K4 methylation was found either globally or at local genes in Mll1-deficient brain cells [20], suggesting that other activities of Mll1 underlie its role in neurogenesis. Thus, the function of H3K4 methylation in NSC self-renewal and lineage determination remains unknown.

We have previously established a direct role of the Dpy30 subunit of the Set1/Mll complexes in facilitating genome-wide H3K4 methylation [21], and also used an unbiased biochemical approach to show that Set1/Mll complex components are the major proteins associated with Dpy30 [14]. These results allow us to investigate the function of efficient H3K4 methylation in various biological processes by perturbing Dpy30’s activity. Interestingly, Dpy30 and efficient H3K4 methylation are not essential for the self-renewal of embryonic stem cells (ESCs), but required for efficient retinoic acid-mediated differentiation of ESCs to neurons and induction of numerous neuronal lineage genes during differentiation [21]. H3K4 methylation conveys plasticity of the transcriptional program during stem cell fate transition [21], and this is further supported by its requirement in efficient reprogramming of differentiated fibroblasts back to the pluripotent state [22]. To understand a role of Dpy30 and efficient H3K4 methylation in tissue-specific stem cells, we generated a *Dpy30* conditional knockout (KO) mouse [23]. Using this mouse model, we have shown a critical requirement of Dpy30 in the correct differentiation as well as long-term maintenance of hematopoietic stem cells [23]. Herein, using *in vitro* and *in vivo* approaches, we sought to determine if Dpy30 and thus H3K4 methylation is involved in self-renewal and differentiation of postnatal NSCs and neural progenitor cells (NPCs, together NSPCs).

## MATERIALS AND METHODS

### Animals

All animal procedures were approved by the Institutional Animal Care and Use Committee at the University of Alabama at Birmingham. All mice were maintained under specific pathogen free conditions and housed in individually ventilated cages. *hGFAP-Cre* mouse (JAX #004600) and *Nestin-Cre* mouse (JAX #003771) were obtained from the Jackson Laboratory. *Dpy30*^*+/-*^ and *Dpy30*^*F/F*^ mice were previously generated in our laboratory [23]. *hGFAP-Cre* mice were crossed with *Dpy30*^*+/-*^ mice to obtain *hGFAP-Cre; Dpy30*^*+/-*^ mice. The latter were then crossed with *Dpy30*^*F/F*^ mice to obtain *hGFAP-Cre; Dpy30*^*F/+*^ (Control) and *hGFAP-Cre; Dpy30*^*F/-*^ (KO) from the same litter. KO mice die naturally around 20-27 days after birth and thus the brains were harvested before 15 days old. Both male and female mice were utilized.

### Tissue preparation, immunofluorescence staining, imaging, and quantification

Tissue was harvested and placed in 4% paraformaldehyde for 48 hours at 4°C. Serial 40 μm, free-floating coronal sections from different bregma levels (together representing 1/6 of the brain) were generated for each immunostaining. These sections were permeabilized in Tris-buffered saline containing TritonX-100 (TBST; 50mM Tris with 0.9% NaCl, and 0.5% TritonX-100) for 10 minutes, incubated with 0.3% H_2_O_2_ for 10 minutes, and blocked with 10% goat serum/TBST for 30 minutes at room temperature. Primary antibodies were incubated in 1% goat serum/TBST for 48 hours rocking at room temperature. Primary antibodies for Dpy30 [21], H3K4me3 (1:500, MP 07-473, EMD Millipore, Billerica, MA), Ki-67 (1:500, Abcam Cambridge, MA ab15580), Blbp (1:300, EMD Millipore, Billerica, MA ABN14), Dcx (1:200, Abcam ab18723), Gfap (1:500, Cell Signaling, Danvers, MA, 3670S), NeuN (1:300, Millipore ABN78), and β-Tubulin III (Tuj1, 1:1000, Biolegend, San Diego, CA 802001) were used. All primary antibodies are of rabbit host except Gfap, which was of mouse host. Secondary antibodies conjugated to Alexa 488 or 546, or 568 (Biolegend, San Diego, CA) were used. The sections were incubated in 1% goat serum/TBST for 4 hours each sequentially in 1:500 dilution of secondary antibodies. Nuclei were labeled with 4’,6-diamidino-2-phenylindole (DAPI, Life Technologies) and mounted in Fluoro Care Anti-Fade mounting media (Biocare Medical, Cat# FP001G10) on 1% gelatin coated microscope slides (Fisher Scientific, Cat#12-550-15). For certain double immunostaining assays where both primary antibodies are from the same species, a sequential staining was performed with excessive washing in between. For Nissl Stain, after mounting free-floating sections on gelatin-coated slides, sections were incubated with 1% cresyl violet acetate, and washed with dH_2_O followed by dehydrated with graded ethanol and xylenes.

All images were collected with an Olympus BX53 fluorescent (Center Valley, PA) microscope with DP72 camera. For *in vivo* cell population and proliferation quantification, the average total number of indicated staining cells were manually counted from 3 different sections for DG and SVZ each from one mouse. This was done for a total of 3 different mice for control and KO each. For quantification of all staining in DG, only cells in focus (on one plane) from both arms of DG were counted. For Dpy30 and H3K4me3, cells with a positive nuclear staining were counted. For Blbp quantification, cells with nuclear staining along with an axon-like projection were counted. For Dcx quantification, cells showing a reticulate/halo of the staining with a long process were counted. For NeuN quantification, positive nuclear staining was counted. For quantification of all SVZ staining except Gfap and Ki-67, total area and area of positive staining was measured using ImageJ (NIH image analysis software, https://imagej.nih.gov/) ‘measure tool’ that provides user with an ‘area count’ in arbitrary units. The area of positive staining was then divided by total area of SVZ to report the ratio. For Gfap quantification, the cells right above the DG and SVZ structure showing radial tubular projection staining were counted. For Ki-67 quantification in both DG and SVZ, red dots within the DG and SVZ were counted. No NeuN was found in the SVZ of control or KO mice.

### Western blotting

Whole fractionated brain cells were lysed by SDS loading buffer (2% SDS, 0.1% Bromophenol blue, 10% glycerol, 100mM DTT, and 50 mM Tris-Cl, pH 6.8) and boiled for 7 minutes. Proteins were resolved on SDS-PAGE gel followed by Western blotting using previously described antibodies [21] for H3, H3K4me1, and H3K4me2, or H3K4me3 (same as used for immunostaining), and Gapdh (EMD Millipore MAB374). Key signals were quantified using ImageJ program (https://imagej.nih.gov/) followed by subtraction of a blank area and normalization to the indicated reference signal intensity.

### Neurosphere Assays

#### Isolation

Progenitors were isolated from 12 days old SVZ tissue using 0.05% trypsin. Neurospheres were grown on non-tissue culture treated plates with mouse NeuroCult proliferation media (Stem Cell Technologies; Seattle, WA) containing 10 ng/ml bFGF, (ProSpec, East Brunswick, NJ), 10 ng/ml EGF (ProSpec), and 2 μg/ml heparin (Fisher Scientific). In the case of progenitors isolated from 12 days old hippocampi, cells from KO mice brain did not grow at all in culture and hence could not be used in serial re-plating or differentiation assays described below.

#### Serial re-plating assay

The process of the directly isolated progenitors (as explained above) in proliferation media for 7 days is defined as “passage 0”. Neurospheres were dissociated using Accutase (Fisher Scientific). Single cell dissociation was plated in 6-well (non-tissue culture treated) plates at 20,000 cells/well in 2 ml of complete proliferation media (thus 10 cells/µl). 7 days later (at passage 1) primary neurosphere number per well was counted and their diameters were measured. Following evaluation, primary neurospheres were collected, dissociated, and re-plated at 10,000 cells/well (thus 5 cells/µl). Secondary sphere number and size were evaluated 7 days later (at passage 2). Secondary spheres were again dissociated and re-plated at 5,000 cells/well (thus 2.5 cells/µl) for 3 more passages to further evaluate self-renewal. Images of neurospheres were taken on a Nikon Eclipse Ti-S inverted microscope system and the diameter of each sphere was measured manually from images using a ruler (and scale bar from images for conversion 2.5 cm = 100 μm for images taken with a 10x objective). Neurospheres at passage 0 were not characterized because of concern of non-clonality and variation in the number of the initiating cells due to variation in the amount of dissected tissue out of control and KO brain. Counting neurospheres for passages 1 and 2 was performed by sampling 100 μl of culture under microscope. The number of neurospheres counted in 100 μl of culture was then multiplied by 20 to get an estimation of total neurospheres in 2ml media. Counting for passages 3-5 was performed from images since the neurospheres then became fewer and could be captured by multiple non-overlapping images covering the whole area of a well.

#### Differentiation assay

Neurospheres at passage 3 were dissociated and plated at 10,000 cells/well in Poly-L-ornithine (Sigma, P3655) and Laminin (Millipore CC095) coated plates in differentiation media (NeuroCult media without growth factors). Staining was performed as described for coronal sections in immunofluorescence protocol at 10 days in differentiation media. Images were taken as described in Serial re-plating assay.

### NPC culture, gene knockdown (KD), growth, differentiation, and chromatin immunoprecipitation (ChIP) assays

For stable KD, ReNcell VM (Millipore, SCC008) and NE-4C (ATCC, CRL-2925) cells were grown and maintained as recommended by the vendor’s protocol. ReNcell VM is an immortalized human neuronal progenitor cell line derived from ventral mesencephalon of human fetal brain and has the ability to readily differentiate into neurons and glia cells. NE-4C is a neuroepithelial cell line derived from cerebral vesicles of 9-day old mouse *p53*^*-/-*^ embryos that differentiate into neurons upon induction with retinoic acid [24]. These cells were infected with lentiviruses expressing control or human or mouse Dpy30 shRNAs [all sequences are available in [25]], or human ASH2L shRNA (5’-CCTGCTTGTATGAACGGGTTT-3’), followed by selection using 2 μg/ml of puromycin (Invivogen, 58-58-2) for 2–3 days starting from 2 days after infection. Live cells were counted using Trypan blue (Gibco, 15250061) exclusion. Cells were then used for RT-qPCR analysis, ChIP-qPCR analysis, and differentiation assays.

For NPC growth assays, puromycin-selected cells were plated at 10,000 cells/ml in the cell maintenance media in the absence of selecting antibiotic to exclude the possibility of altered expression of antibiotic-resistance gene, and number of live cells was counted using a hemocytometer on indicated days. For NPC differentiation assays, puromycin selected cells were plated at 100,000 cells/well in a 12-well plate and maintained at 1.2 μg/ml puromycin. ReNcell VM cells were cultured in differentiation media that consists of cell maintenance media with puromycin, but without growth factors. NE-4C cells were differentiated by adding 1 μM all-trans retinoic acid (Sigma, 302-79-4) to the culture media with puromycin. Nine days into differentiation, cells were counted, harvested for RNA isolation and measurement of mRNA levels, and stained as described for coronal section staining in Immunofluorescence protocol. Quantification was performed on images taken from 3 independent assays. In dual staining assays, completely overlapping yellow dots were not counted. Partially overlapping yellow dots were counted as 1 for neuron and 1 for glia since these are different cells growing partially on top of each other.

NPC ChIP assays were performed as previously described [21]. For quantifications of ChIP results, percent input was first calculated from ChIP qPCR data as previously described [21], and the relative enrichment fold was calculated as the ratio of the percent input value for each locus over that for the *Map2* or *MAP2* locus in the scramble control cells (which was set to 1).

### qPCR, RNA-seq and bioinformatics

Total RNAs were extracted from freshly isolated (micro-dissected) SVZ tissue or hippocampi using Direct-zol™ RNA MiniPrep kit (Zymo Research). Some RNAs were reverse transcribed with SuperScript III (Invitrogen). qPCR was performed with SYBR Advantage qPCR Premix (Takara Bio Inc.) on a ViiA7 Real-Time PCR System (Applied Biosystems). Primers used are listed in Table S3. Relative expression levels were normalized to indicated genes in the figure legends. Total RNAs were sent to the Genomic Services Lab at the HudsonAlpha Institute for Biotechnology (Huntsville, AL) for sequencing, including library construction. In brief, the quality of the total RNA and DNA was assessed using the Agilent 2100 Bioanalyzer. Two rounds of polyA+ selection was performed for RNA samples, followed by conversion to cDNAs. We used the mRNA library generation kits (Agilent) and TruSeq ChIP Sample Prep kit (Illumina) per the manufacturer’s instructions. The indexed DNA libraries were quantitated using qPCR in a Roche LightCycler 480 with the Kapa Biosystems kit for library quantitation (Kapa Biosystems) before cluster generation. Clusters were generated to yield ∼725–825 K clusters/mm^2^. Cluster density and quality were determined during the run after the first base addition parameters were assessed. Sequencing was performed on Illumina HiSeq2500 with sequencing reagents and flow cells providing up to 300 Gb of sequence information per flow cell. For RNA-seq, we obtained 35– 55 million reads 51-bp paired-end reads for each RNA sample. Data analysis was performed using TruSeq Stranded RNA-Seq Pipeline in Partek Flow with some parameter modifications. Briefly, low quality mapped reads (MQ < 30) were removed from the analysis, mapped to the mouse reference genome (mm10) using TopHat2 (v2.1.0), and annotated using Gencode v20 (changed parameter “Strand Specificity” set to “Auto detect”). Read counts were normalized by TMM then FPKM and differential gene expression analyses (GSA) were performed to identify genes differentially expressed between control and KO tissues. Gene ontology analysis was performed with DAVID 6.7 (https://david-d.ncifcrf.gov/). GSEA was performed at http://software.broadinstitute.org/gsea/index.jsp. The gene sets were obtained from previously deposited and curated data in mSigDBv3.0 database (http://software.broadinstitute.org/gsea/msigdb). All gene sets used with the exception of ‘Neuroblast migration’ were previously deposited in Molecular Signatures Database at http://software.broadinstitute.org/gsea/msigdb/index.jsp. Gene sets involved in neuroblast migration were curated from [26-31]. Gene sets enriched in quiescent NSC population or activated NSC population were from [32]. Genes were rank-ordered in descending order according to the fold change. The list of pre-ranked genes was analyzed with the gene set matrix composed file (.gmx) from curated data. Significant gene sets enriched by Dpy30 KD were identified using nominal P value < 0.05 and FDR value < 0.1. All analyses were performed using GSEA v2.0 software with pre-ranked list and 1000 data permutations.

Our RNA-seq datasets have been deposited in the Gene Expression Omnibus database under GSE114091.

### Statistics

All data are reported as mean ± SD. Unless indicated otherwise in legends, unpaired and equal variance, two-tailed Student’s t test was used to calculate P-values and evaluate the statistical significance of the difference between control and KO samples in all comparisons. F-test was used to determine if variances are equal. P value less than 0.05 was considered significant. For Figure 3D left and middle panels, a Mann-Whitney *U* test was performed to evaluate the difference between control and KO neurosphere number and subsequent drop between consecutive passages at alpha 0.05. For Figures 4B, 4F, and S4C, all samples comparison was performed using Two-factor ANOVA with replication, followed by post hoc t-Test. *P < 0.05, **P < 0.01, ***P < 0.001, ****P < 0.0001, and *****P < 0.00001.

### Video

Video was recorded using iPhone 6 camera.

## RESULTS

### Expression of Dpy30 in the developing brain

To learn the expression pattern of Dpy30 in the developing mammalian brain, we mined a collection of *in situ* hybridization images for gene expression in the developing and adult mouse nervous system [33]. We found that Dpy30 is mainly expressed in the hippocampus and the cerebellum starting at E15 but it’s expression is diminished by early adulthood (P42) (Figure S1A). These results suggest a possible role of Dpy30 during a specific window of brain development. To determine if Dpy30 is expressed in NSCs, NPCs, and their progeny, we mined a recently published single-cell RNA-sequencing (RNA-seq) dataset of these cells in the mouse neocortex [34], and found Dpy30 is expressed at the highest level in NSCs and intermediate progenitor cells but low levels in early or late neurons (Figure S1B). We also examined the expression of the other core subunits and the catalytic subunits of the Set1/Mll complexes. Interestingly, all of the other core subunits, including Ash2l, Rbbp5, and Wdr5 are expressed at low levels in NSCs, NPCs, and neurons. Among the six catalytic subunits in a Set1/Mll complex, Kmt2a/Mll1, Kmt2c/Mll3, and Kmt2d/Mll4 are expressed at high levels in NSCs, NPCs, and neurons, while Kmt2b/Mll2, Setd1a, and Setd1b are expressed at low levels in these cell types (Figure S1B). Dpy30 is thus the only subunit in the Set1/Mll complexes that is preferentially expressed at a high level in NSCs and NPCs, again suggesting an important role of Dpy30 in early stages of neural development.

To further investigate the specific cell types that express Dpy30, we performed immunofluorescence staining of Dpy30 and neuronal nuclei (NeuN), a nuclear marker protein for mature neurons. In the hippocampal dentate gyrus (DG), we saw expression of Dpy30 in both mature granular neurons and cells within the subgranular zone. Dpy30 was also expressed in cells within the subventricular zone (SVZ) and adjacent neuronal layers (Figure S1C). As the DG and SVZ are the major areas with significant neurogenesis into adulthood [1], expression of Dpy30 here prompted us to pursue a functional study of Dpy30 in mammalian brain development and postnatal neurogenesis.

### Dpy30 deficiency results in reduction of H3K4 methylation and failure of postnatal neurogenesis

To study the biological function of Dpy30 in postnatal neurogenesis, we used the human *glial fibrillary protein (hGFAP)-Cre* [35] *to excise the floxed Dpy30* allele in NSCs present at embryonic day (E) 13.5 in the dentate, cerebellar granular cell layer, and SVZ [36]. To avoid potential effects from *Cre* expression, we restricted our comparisons between *hGFAP-Cre; Dpy30*^*F/+*^ and *hGFAP-Cre; Dpy30*^*F/-*^ littermates as confirmed by genotyping (Figure S1D), hereafter referred to as control and KO, respectively. The KO mice were born at the expected Mendelian ratios and were indistinguishable from their control littermates. However, by postnatal day (P) 5, the KO mice developed progressive but severe growth retardation (Figures S1E and S1F) and ataxia (video S1), and died between P20 and P27. This phenotype is 100% penetrant in a total of over 200 offspring animals. Moreover, we also deleted *Dpy30* using *Nestin*-*Cre* [37], whose expression starts E12.5 in cortical prepalate layer and increases during perinatal development in NSCs of the subependymal zone and in NPCs of the rostral migratory stream of newborn mice [38]. The *Nestin-Cre; Dpy30*^*F/-*^ mice exhibited essentially identical phenotypes of growth retardation (Figure S1G) and ataxia as the *hGFAP-Cre; Dpy30*^*F/-*^ mice, further corroborating a requirement of Dpy30 in postnatal brain development. We have thus only used the *hGFAP-Cre; Dpy30*^*F/+*^ (control) and *hGFAP-Cre; Dpy30*^*F/-*^ (KO) littermates for the rest of this study.

While most of the brain appeared morphologically normal, the hippocampal DG showed grossly altered ultrastructure and the lateral ventricles were enlarged (Figure 1A). Moreover, KO mice show a marked reduction in cerebellar folia (marked with the arrows in Figure S1H) and cells in the internal granular layer (IGL) (Figure S1I). *Dpy30* mRNA levels were significantly reduced in the KO SVZ and DG (Figure S1J). As shown by immunostaining, the Dpy30 protein was almost completely lost in the KO SVZ, DG, and cerebellar granule cell layer (Figure 1B-1D). These results confirmed that the *hGFAP-Cre; Dpy30*^*F/-*^ mice were deficient of Dpy30 in the specific regions that also show gross morphological changes. Consistent with a role of Dpy30 in regulating all three levels of H3K4 methylation [21], we detected a modest reduction of H3K4me1, and a significant reduction of H3K4me2 and H3K4me3, by Western blotting of whole brain KO lysates (Figure 1E). Confirming this result visually, immunohistochemistry shows a stark decline in H3K4me3 staining in the DG, SVZ, as well as the cerebellar granule cells of the KO brain, compared to the control (Figure 1B-1D). As these are the major brain regions that undergoes substantial postnatal neurogenesis, they overall suggest an important role of Dpy30 and efficient H3K4 methylation in postnatal neurogenesis.

**Figure 1.**
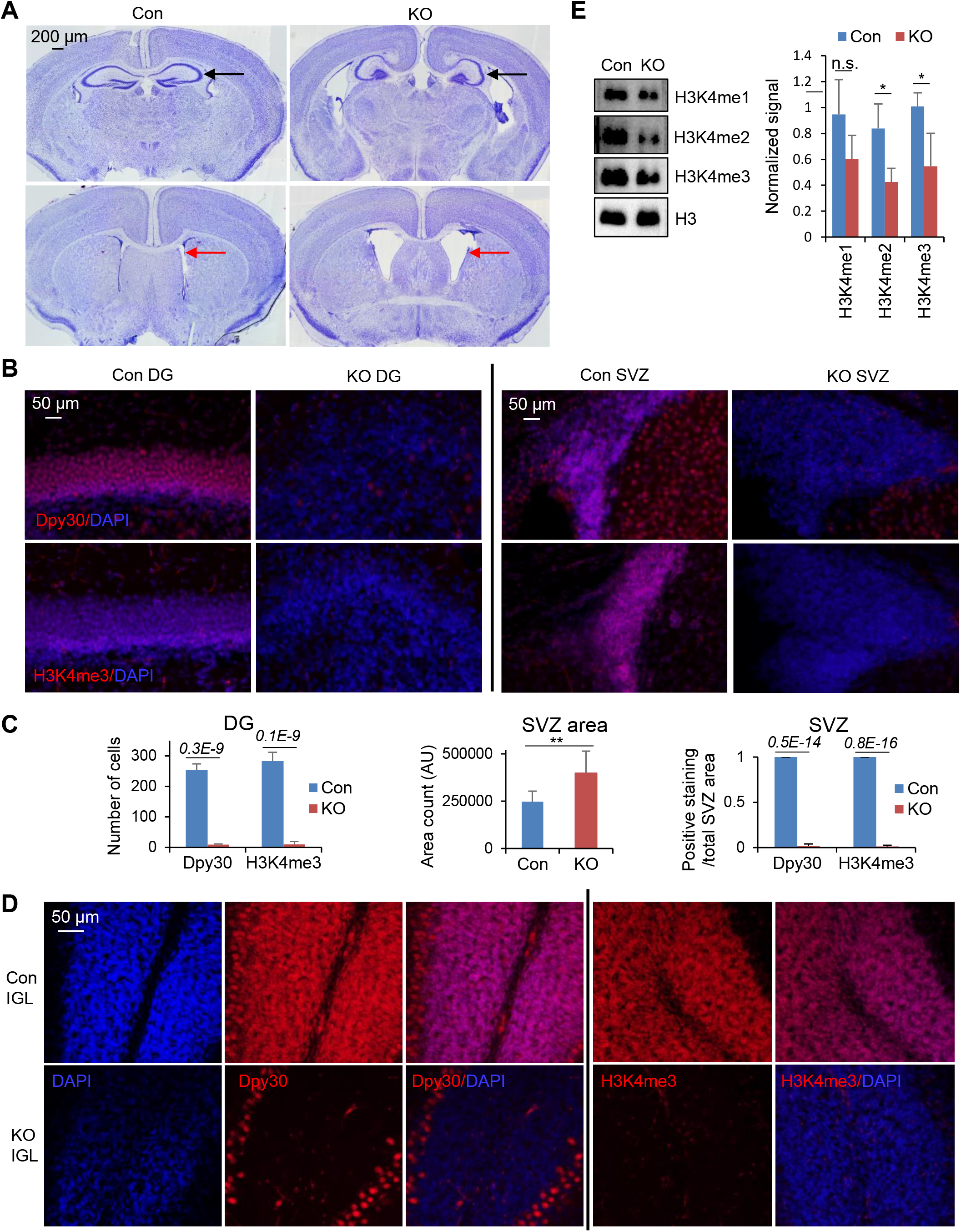
Dpy30 deficiency results in defective postnatal brain development. **(A)** Nissl staining of coronal sections from P12 control (Con) and KO mice. Representative of 3 mice each. Black arrows point to hippocampus and red arrows point to SVZ. **(B)** Staining for DAPI and Dpy30 (top) or H3K4me3 (Bottom) in coronal sections from P12 control and KO mice. Representative of 3 mice each genotype. **(C)** Quantification of staining in (B). **(D)** Staining for DAPI and Dpy30 (left) or H3K4me3 (right) in coronal cerebellum sections from P12 control and KO mice. Representative of 3 mice each genotype. IGL, Internal granular layer. **(E)** H3K4 methylation levels in the whole brain lysates from P12 control and KO mice were determined by Western blotting. Shown are representative results (left) and quantifications for the band intensity relative to the total H3 band intensity of the same sample (right). n = 4 mice each genotype. Data represent the mean ± SD for (C and E). P values are either labeled on top of the bars or \**P* < 0.05 and \*\**P* < 0.01, by two-tailed Student’s t test.

### Transcriptome analyses suggest a requirement of Dpy30 for NSPC fate determination

To begin to understand the alterations caused by Dpy30 loss at the molecular level, we examined the gene expression profiles of micro-dissected hippocampal (containing DG) and SVZ tissues extracted from 3 different P12 mice of each genotype via RNA sequencing (RNA-Seq) (Tables S1 and S2). Our Gene Ontology (GO) analysis showed that the top 10 most significantly over-represented gene functions downregulated in the KO hippocampus were involved in neuronal activities, such as synapse and ion channel activities (Figure S2A, top left). More specific analyses by Gene Set Enrichment Analysis (GSEA) [39] also revealed significant enrichment of neuronal markers [40] among genes downregulated in the KO hippocampus (Figure 2A, left), which was confirmed by qPCR for the top group of neuronal marker genes (Figure S2B and S2C). In addition, some neuronal genes that are not included in the gene set for neuronal markers were also downregulated by Dpy30 loss in the hippocampus. For example, Glutamate Ionotropic Receptor NMDA Type Subunit 2C (*Grin2c*), a neuronal specific gene [41], was among the top 50 downregulated genes in KO hippocampus (Figure S2E, left) and also shown by qPCR to be largely downregulated (Figure S2C). Enrichment (but not statistically significant) of astrocytic markers [40] was also observed in genes downregulated in the KO hippocampus (Figure S2D). These results suggest a relatively selective loss of neurogenesis and/or neuronal function in the Dpy30-deficient hippocampus.

**Figure 2.**
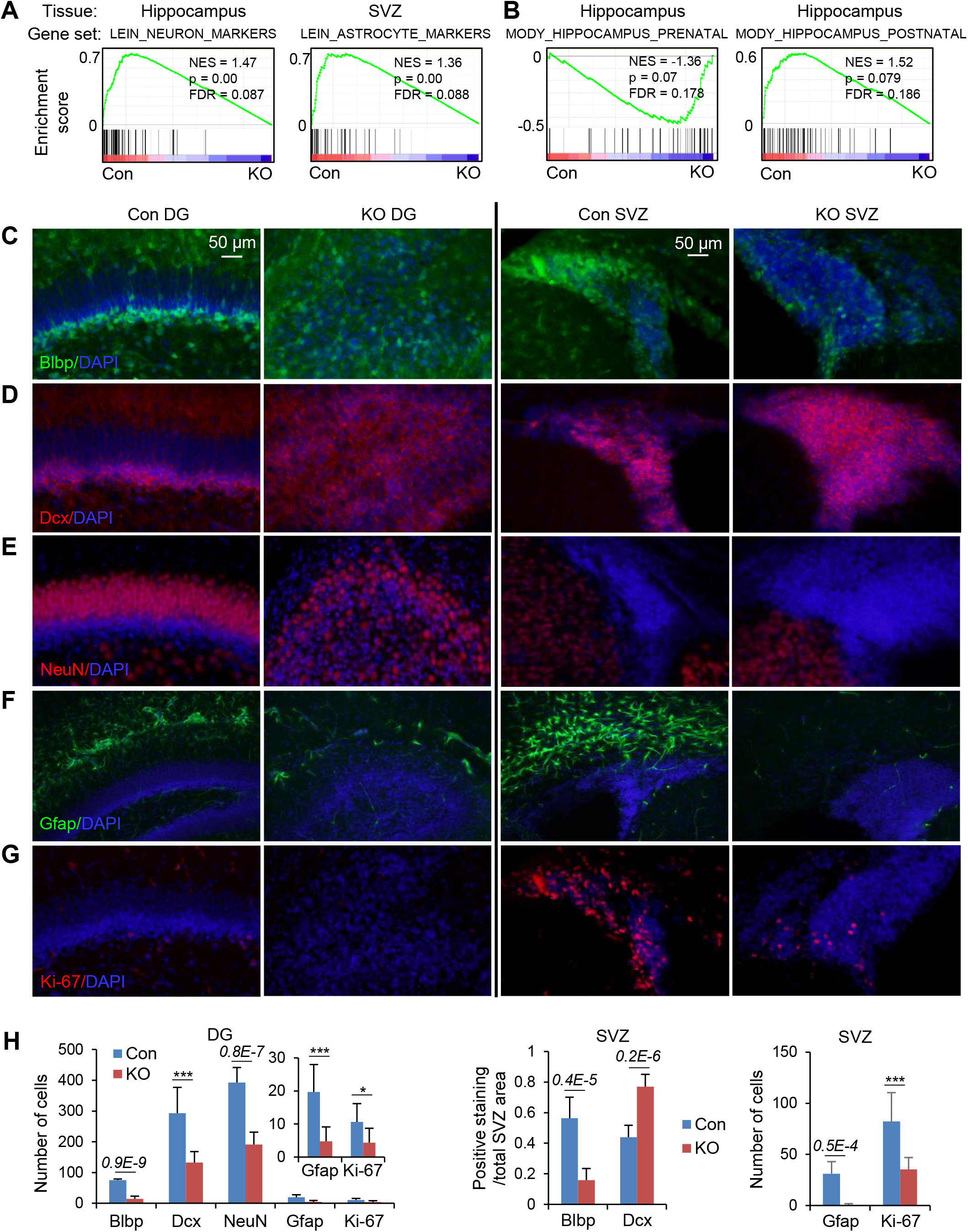
Dpy30 deficiency impairs neurogenesis and gliogenesis in the postnatal brain. **(A-B)** GSEA for RNA-seq results from Hippocampus or SVZ cells (indicated) of P12 control and KO mice (n = 3 mice each) using indicated gene sets. NES, normalized enrichment score. **(C-G)** Staining for DAPI and Blbp (C), Dcx (D), NeuN (E), Gfap (F), Ki-67 (G) in coronal sections from P12 control and KO mice. Representative of 3 mice each genotype. **(H)** Quantifications of positive stained cells or area of the images in C-G. Both top and bottom Subgranularzone/granular zone layers of the dentate were counted for control and the whole dentate (absence of a subgranualr zone) was counted for KO. For SVZ, ImageJ software’s “area count” was used to obtain area of positive staining and total area of SVZ. n = 9 (3 mice, each with 3 different bregma levels). Data represent the mean ± SD in (H). P values are either labeled on top of the bars or \**P* < 0.05, \*\**P* < 0.01, and \*\*\**P* < 0.001, by two-tailed Student’s t test.

On the other hand, the most enriched gene functions downregulated in the KO SVZ were not in mature neuronal activities, but rather included mostly GTPase regulator activities (Figure S2A, bottom left), which are known to regulate migration of neuronal precursors and neurite outgrowth [42, 43]. Immature neurons/neuroblasts within SVZ travel from SVZ to the olfactory bulb via the rostral migratory stream and differentiate during that process [44-46]. Indeed, GSEA showed that genes associated with neuroblast migration were significantly enriched among downregulated genes in the KO SVZ, but not in the KO hippocampus (Figure S2D). These results suggest the possibility of defective precursor migration from SVZ to the olfactory bulb for differentiation. Different from the downregulation of neuronal genes in the KO hippocampus, GSEA showed that astrocytic markers [40] were significantly enriched in genes downregulated in the KO SVZ (Figure 2A, right), which was confirmed by qPCR for the top group of astrocytic marker genes (Figure S2B and S2C). Once again, some astrocytic marker genes not included in the gene set were also downregulated in the KO SVZ. For example, Solute Carrier Family 15 Member (*Slc15a2*), a gene selectively expressed in astrocytes [47-49], was among the top 50 most downregulated genes in the KO SVZ (Figure S2E, right). *S100B* is another gene that is also specifically expressed in astrocytes [50]. Our qPCR assays confirmed the significant downregulation of both *Slc15a2* and *S100B* in the KO SVZ (Figure S2C). We found no significant change in neuronal markers upon loss of Dpy30 in SVZ (Figure S2D). These results thus suggest a relatively specific disruption of gliogenesis in the Dpy30-deficient SVZ.

Interestingly, the top enriched gene functions upregulated upon Dpy30 loss in the KO hippocampus and those in the KO SVZ both included mostly ribosomal genes (Figure S2A, right two panels), suggesting an increase in protein synthesis activity in both of these NSC-related tissues. Consistent with this finding, GSEA also revealed upregulation of activated NSC gene signatures [32] in the KO hippocampus, including genes associated with protein metabolism, mRNA metabolism, and DNA repair (Figure S2F). On the other hand, GSEA for the KO SVZ showed downregulation of pathways enriched in quiescent stem cells [32], such as lipid metabolism, growth factor activity, and Peroxisome Proliferator-Activated Receptor (PPAR) signaling (Figure S2G). These results suggest that Dpy30 may regulate the pool of quiescent NSCs in the major postnatal neurogenic regions in brain.

By performing GSEA using a gene set characterized for genes upregulated at prenatal, neonatal, and postnatal developmental stages of the mouse hippocampus [51], we found enrichment of genes characteristic for prenatal hippocampus in those upregulated in the KO hippocampus, as well as enrichment of genes for postnatal hippocampus in those downregulated the KO hippocampus (Figure 2B). For example, Aldolase, Fructose-Bisphosphate C (*Aldoc*), a gene selectively expressed in the postnatal hippocampus [52], was among the top 50 downregulated genes in KO hippocampus (Figure S2E, left) and its significant downregulation was also shown by qPCR (Figure S2C). In other words, the gene expression program of the P12 Dpy30-deficient hippocampus resembled a prenatal stage rather than a normal postnatal stage, strongly suggesting a developmental arrest of the Dpy30-deficient hippocampus. Finally, while a NSC-specific signature gene set is not readily available for GSEA, our qPCR assays showed that *Prominin1* (*Prom1*) and *Gfap*, the coincident expression of which marks adult NSCs *in vivo* [53], *were both significantly downregulated in the KO SVZ (Figure S2C). Although our assays do not provide information on co-expression of both markers in the same cell and Gfap* downregulation is more likely a result of reduced gliogenesis, this is consistent with a reduced NSC pool or activity in the KO SVZ.

### Dpy30 deficiency impairs neurogenesis and gliogenesis in the postnatal brain

The transcriptome data prompted us to further investigate the effect of Dpy30 loss on NSC activity in the developing brain, especially DG and SVZ. NSCs present in both of these regions express Brain lipid binding protein (Blbp). Consistent with the downregulation of NSC marker *Prom1* in the KO SVZ, we found that the Blbp-expressing cell population was significantly reduced in both DG and SVZ of the KO mouse, compared to their corresponding tissues from the control mice (Figure 2C and 2H). DG of KO mice exhibited reduced immature and mature neuronal populations as seen by doublecortin (Dcx) and NeuN staining, respectively (Figure 2D and 2E, left, and 2H). Most striking, however, is the extreme disruption of dentate morphology for both cell groups where it is impossible to distinguish distinct granular and subgranular zones. This is consistent with the reduction of neuronal markers in the KO hippocampus shown by the transcriptome analyses. DG also showed a significant decrease in Gfap-expressing astrocyte population (Figure 2F, left). Due to differences in the cellular composition around the two neurogenic regions of the postnatal brain [54], SVZ and DG appeared to respond differently to the excision of *Dpy30*. P12 KO SVZ exhibited an accumulation of immature neurons as seen by increased Dcx staining (Figure 2D, right, and 2H, middle). the significant downregulation of genes associated with neuroblast migration shown by the transcriptome analyses above, these results are most consistent with a disruption of neuroblast differentiation and/or migration, leading to immature neuron accumulation in the KO SVZ. Additionally, the KO SVZ showed a drastic reduction in the astrocyte population as seen by staining for Gfap (Figure 2F, right, and 2H, right), consistent with the enrichment of astrocytic markers among genes downregulated in the KO SVZ.

Finally, we found that, compared to their corresponding control tissues, both the DG and SVZ from the KO mice had significantly less actively dividing cells expressing proliferation protein Ki-67 (Figures 2G and 2H) [55] that is also functionally crucial for cell division by maintaining the structural integrity of mitotic chromosomes [56]. Therefore, Dpy30 is required to generate or maintain the pool of proliferative NSCs.

### Dpy30 is required for sustaining self-renewal and proliferation of NSPCs *in vitro*

To determine whether Dpy30 is cell-intrinsically required for self-renewal of NSCs, we extracted NSCs from SVZ of KO mice, and measured their self-renewal using the neurosphere serial re-plating assay (Figure 3A) [12, 57]. We first confirmed our technical ability to isolate and grow multipotent control neurospheres. Control neurospheres differentiated into both neuronal and glial lineages with characteristic cell morphologies and lineage markers upon appropriate treatment (Figure 3B and S3A). However, KO neurospheres show low viability which precluded detailed analysis of neurospheres. We did note that serial re-plating assays of the KO cells showed a significant reduction in both the number and size of neurospheres compared to the control cells in each of the same passages (Figure 3C and 3D), and unlike controls, there were no live cells in the KO culture after the 5th passage (Figure 3C and 3D). Experiments to model self-renewal in hippocampal DG-derived NSCs were not possible as the cultures of the KO hippocampal tissues produced no viable cells (Figure S3B). These results indicate that Dpy30 is required for efficient self-renewal of NSCs and/or the proliferation of NPCs in a cell-autonomous manner. In particular, the progressive increase in clonogenicity loss of the KO SVZ cells along each round of re-plating (Figure 3D, middle) argues that the effect was not merely an accumulated result of Dpy30 loss since the embryonic stage, but rather an intrinsic defect in self-renewal as a result of Dpy30 loss in the assayed cells.

**Figure 3.**
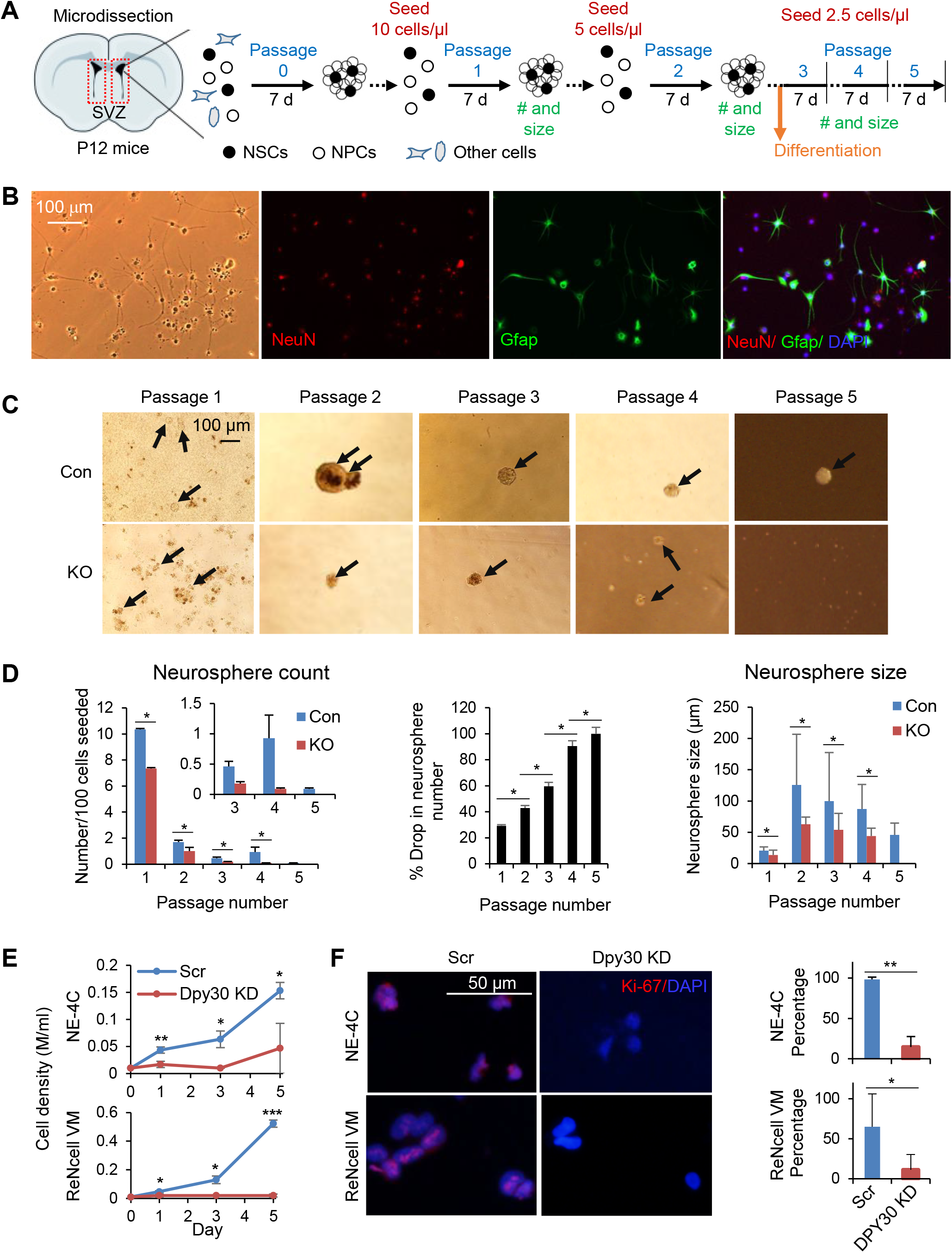
Dpy30 is required for sustaining the self-renewal and proliferation of NSPCs. **(A)** Diagram for the neurosphere assays including serial re-plating and differentiation. **(B) I**mages of differentiated neurospheres obtained from SVZ of control mice. NeuN for neurons and Gfap for glia. Representative of 3 control mice. **(C)** Serial re-plating assay with NSCs isolated from SVZ of control and KO mice. Representative of 2 mice each. Images were taken 7 days into the indicated passage (top) except passages 1 and 4, which were taken 5 days into the passage. **(D)** Quantifications of number of neurospheres per 100 cells seeded (left), percent drop in neurosphere number (middle), and neurosphere size (right) at each passage. n = 2 mice each genotype. Percent drop was calculated as (control count - KO count)/control count x 100% at each passage. Neurosphere size was calculated by measuring the diameters of multiple neurospheres from 2 mice each genotype. Total measured neurosphere number n = 50 for each genotype at passage 1, n = 39 (control) or 6 (KO) at passage 2, n = 46 (control) or 18 (KO) at passage 3, n = 93 (control) or 8 (KO) at passage 4, and n = 9 (control) at passage 5. There was no KO neurosphere in passage 5. **(E and F)** Growth assays (E) and staining for Ki-67 and DAPI (F) for stably selected control (Scr) and Dpy30-KD NE-4C mouse NPCs (top) and ReNcell VM human NPCs (bottom). Shown in (F) are representative images (left) and quantifications (right). n = 3 independent infections for all. Data represent the mean ± SD for (D-F). \**P* < 0.05. \*\**P* < 0.01. \*\*\**P* <0.001, by Mann-Whitney *U* test (alpha = 0.05) for middle and left panels in (D), and two-tailed Student’s t test for (D) right panel, (E), and (F).

To test the broader requirement of Dpy30 activity for NSPC activity and further address if the effects on neurogenesis only reflect an accumulated impact of Dpy30 loss, we sought to use shRNAs to acutely deplete Dpy30 from NE-4C, a mouse neural progenitor cell (NPC) line, and ReNcell VM, a human NPC line in culture. Knockdown (KD) of Dpy30 in these cells drastically decreased their ability to grow in culture (Figure 3E). Moreover, percentage of the Ki-67-positive cells was also significantly reduced by Dpy30 KD in these cells (Figure 3F). This suggests that Dpy30 is required for maintaining the NSPC proliferation.

### Dpy30 directly regulates differentiation of NPCs to neuronal and glial lineages

We next studied the effect of Dpy30 KD on the differentiation of these cultured NPC lines. Upon treatment of differentiation conditions, while the control mouse NPCs differentiated into cells positive for β-Tubulin III (Tuj1) staining with extensive fiber-containing morphology that represent neuronal axons, the Dpy30-KD mouse NPCs failed to produce any cells that are β-Tubulin III-positive with fiber-like structure (Figures 4A and S4A). We were unable to determine the effect on their glial differentiation as the control mouse NPCs also did not successfully differentiate to glial lineages (data not shown). DPY30 KD in the human NPCs clearly abolished their glial differentiation as shown by the extensive GFAP-positive cells seen after differentiation of the control NPCs but the drastic reduction of the number and percentage of these cells for the DPY30-KD NPCs (Figures 4D, 4E, and S4B). Regarding neuronal differentiation of the human NPCs, while we detected significantly lower number of the NEUN-positive cells for the DPY30-KD compared to the control cells after providing differentiation conditions (Figures 4D and S4B), the percentage of NEUN-positive cells was not significantly different between control and KD due to the lower number of the overall KD cells (Figures 4E and S4B), suggesting that NeuN expression per se may not be critically dependent on Dpy30. However, even with the greatly underestimated quantification of the fibers in the differentiated control cells (due to their extensively intertwined structures), we found that the KD cells gave rise to significantly lower percentage of cells with long fibers/processes that are characteristic of neuronal axons (Figures 4E and S4B, bottom). These results suggest inefficient neuronal differentiation of the human NPCs upon DPY30 KD. We also examined the effect of depletion of ASH2L, the direct interaction partner of DPY30 and also a core subunit of the SET1/MLL complexes [58] in human NPCs. While we were able to achieve approximately 50% KD efficiency before differentiation, the KD efficiency became very low after treatment of differentiation conditions (Figure S4C), suggesting that cells with efficient ASH2L depletion were not compatible with the differentiation conditions and were likely selected against during the process.

**Figure 4.**
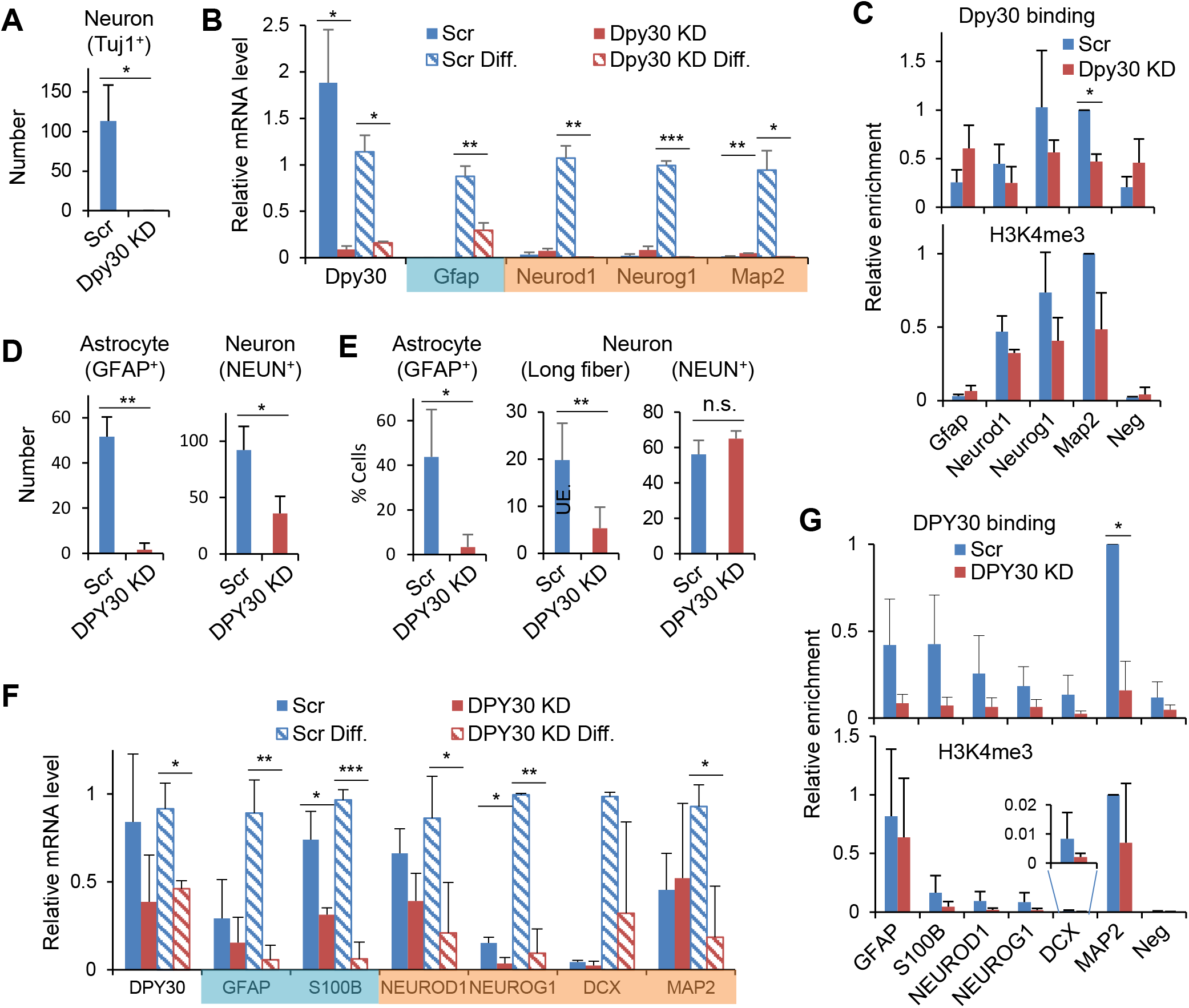
Dpy30 directly regulates NPC differentiation and induction of key genes for differentiation. **(A-B)** Differentiation of NE-4C mouse NPCs, and assayed 9 days into differentiation. n = 3 biological repeats. **(D-F)** Differentiation of ReNcell VM human NPCs, and assayed 9 days into differentiation. n = 3 biological repeats. **(A and D)** Number of neurons and astrocytes (indicated) found in the differentiated control and KD culture. For (A) a percentage plot was not possible due to extensively overlapping DAPI-positive cells in control (Scr) culture. As no Tuj1-positive cells were seen in KD culture, the conclusion based on cell number would be in agreement with percent cell calculations. Scr, scrambled control shRNA. Neurons were identified based on Tuj1 (for mouse) or NEUN (for human) staining, and astrocytes based on GFAP staining. **(E)** Percent astrocytes and neurons found in Scr and DPY30-KD ReNcell VM human NPCs upon differentiation. Astrocytes were identified based on GFAP staining. Neurons were identified based on long (> 20 μm) fibers or NEUN staining. The plots show the percentage of the number of cells with indicated characteristics in the total number of DAPI-positive cells. UE, underestimated, due to extensively intertwined structure. **(B and F)** Relative mRNA levels were determined by qPCR and normalized to *Gapdh/GAPDH* for indicated genes upon *Dpy30/DPY30* KD with and without treatment of differentiation conditions in NE-4C (B) or ReNcell VM (F) cells. One of the biological repeats for ‘Scr Differentiated’ was set to 1 for each gene. Gene names are shaded in aqua for gliogenesis and in orange for neurogenesis. **(C and G)** Dpy30 and H3K4me3 ChIP-qPCR in NE-4C (C) or ReNcell VM (G) cells. Relative enrichment was generated by calculating percent input for each locus followed by normalizing to the percent input for the *Map2/MAP2* locus in the control cells. Data represent the mean ± SD. Two factor ANOVA analysis followed with post hoc *t*-Test was performed in (B) and (F). \**P* < 0.05, \*\**P* < 0.01, and \*\*\**P* < 0.001, by two-tailed Student’s t test for the rest of the figure.

To further understand the differentiation defect of Dpy30-KD mouse and human NPCs, we analyzed the expression change of a number of genes known to regulate NSC or NPC fate determination. These genes include *NEUROD1* [59] *and NEUROG1* [60], which encode E-box-binding transcriptional activators critical for neuronal differentiation, and *MAP2*, which encodes a cytoskeletal protein determining and stabilizing dendritic shape during neuron development [61]. We did not find downregulation of some of these genes in the micro-dissected *Dpy30* KO hippocampal or SVZ tissues (Figure S2C), and suspected that contamination of Dpy30-intact cells may contribute to the lack of downregulation, and thus sought to investigate these genes in the much more homogenous cell lines in culture. These genes were induced after differentiation in the control cells but failed to be induced under the same differentiation conditions in both mouse and human NPCs depleted of Dpy30 (Figure 4B and 4F). We also examined the expression change of genes involved in gliogenesis such as *GFAP* and *S100B*. Similar to the effect on the neurogenic genes, *GFAP* expression was elevated upon differentiation of control NPCs, but such induction was significantly impaired or completely lost in NPCs depleted of Dpy30 (Figure 4B and 4F). The expression of *S100B* was significantly reduced by DPY30 KD before differentiation, and this reduction became more prominent after differentiation of the human NPCs (Figure 4F). Even in the cells that started with decent but ended with minimal depletion of ASH2L under the differentiation conditions, the expression of both the neurogenic and gliogenic genes was modestly impaired compared to the control at the end of the differentiation process (Figure S4C). These results indicate a requirement of the mammalian H3K4 methyltransferase complexes in efficient expression of the lineage specification genes during NPC fate determination.

As shown by Chromatin immunoprecipitation (ChIP) assays, Dpy30 bound to the transcriptional start sites (TSSs) of these genes (Figure 4C and 4G, top, and S4D and S4E, top) in at least one of the NPC models, suggesting that Dpy30 directly controls the expression of these key neurogenesis genes. As a consequence of reduced Dpy30 binding to these genes upon Dpy30 KD, H3K4me3 was reduced at many of these genes (Figure 4C and 4G, bottom, and S4D and S4E, bottom) except *Gfap/GFAP* (see Discussion), although the large variations among biological repeats prevent showing a statistical significance. These results further support a direct and cell-intrinsic role of Dpy30 and its associated H3K4 methylation in the fate determination of neural stem and progenitor cells.

## DISCUSSION

Our results from both *in vivo* and *in vitro* assays strongly suggest a requirement of Dpy30 for the self-renewal and differentiation of NSCs in a cell-intrinsic manner. NSCs in DG and SVZ are significantly reduced upon loss of Dpy30. Cells extracted from the KO DG were unable to form neurospheres *in vitro*. Although the cells from the KO SVZ were able to form neurospheres *in vitro*, the serial replating assays clearly showed a gradually increased loss of neurospheres along each round of replating of the same number of cells from the KO SVZ. Consistent with the reduction of Ki-67-positive cells in the KO DG and SVZ, acute depletion of Dpy30 in human or mouse NPCs greatly reduced their growth and Ki-67 expression, further supporting a requirement of Dpy30 in NSPC proliferation. DPY30 and ASH2L also enable the differentiation capacity of NPCs to both neuronal and glial lineages by directly regulating the epigenetic modifications of key genes in these pathways. This is consistent with a requirement of Dpy30 and Rbbp5 for the neuronal differentiation from ESCs [21], where these H3K4 methylation modulators epigenetically prime the bivalently marked lineage-specific genes for efficient induction upon differentiation cues.

A few lines of evidence support that the phenotypic defects of the Dpy30 KO brain are at least partially due to postnatal loss of Dpy30, including our *in vitro* neurosphere assays that revealed intrinsic defects along serial re-plating of the postnatal brain tissues, and the profound reduction of cells in the internal granular layer (IGL) of cerebellum (Figure S1I), which is known to be postnatally generated. However, considering the high expression of Dpy30 at hippocampus and cerebellum at both the late embryonic and postnatal stages (Figure S1A), Dpy30 most likely regulates both of these stages of neural system development, and the *in vivo* defects are probably a combined result of Dpy30 loss in both of these stages.

Our gene profiling results also support a developmental arrest of the hippocampus upon Dpy30 loss, and begin to reveal molecular pathways governed by Dpy30 and H3K4 methylation in postnatal neuronal and glial development. Due to the technical difficulty of extracting pure cell populations and the high heterogeneity of the tissues in DG and SVZ, our gene profiling results from pooled cells do not necessarily show gene expression changes in NSCs, but most likely also reflect changes of cellular composition, such as increase in metabolically activated cells. However, either interpretation supports a halted development of hippocampus and gliogenesis in SVZ following loss of Dpy30 and H3K4 methylation. Moreover, the unpure cell populations could mask changes of expression of certain genes, such as *Neurod1* and *Neurog1* (Figure S2C), which are directly regulated by Dpy30 for efficient induction in the much more homogenous NPC lines (Figure 4B, 4C, 4F, and 4G). Alternatively, these genes may be robustly regulated *in vivo* through other mechanisms that were not active in the cultured cells. While we have shown the significant change in expression of a number of neurogenesis genes in DG and SVZ of the KO brains, we note that the strong cellular phenotypes may not necessarily arise from significant changes in expression levels of individual genes. Chromatin modulators such as Dpy30 globally (but not uniformly) influence the genome compaction and accessibility, and the collective effects from subtle changes in numerous genes may result in significant cellular phenotypes. Moreover, chromatin modulators can impact gene expression more than just their expression levels, and those effects may not be captured by the current RNA-seq technology. For example, the breadth of H3K4me3 domain has been linked to transcription consistency and cell identity [23, 62], and the top 30 broadest H3K4me3 domains in NSPCs were proposed to be novel regulators of NSPC self-renewal and/or neurogenesis [62], although our analyses here did not show significant enrichment of these genes among genes up- or down-regulated in the KO SVZ and hippocampus (Figure S2H).

Dpy30 was originally discovered as a gene required for dosage compensation and development of *C. elegans*. Dpy30 mutations cause complete lethality in XX *C. elegans* and numerous abnormalities in XO animals including uncoordinated movement [63], suggesting a requirement of Dpy30 for neuromuscular development. Here, we show that loss of Dpy30 in neurogenic regions (including the cerebellar cells) of the brain results in ataxia like uncoordinated movement in these mice. The deficiency of Dpy30 and H3K4 methylation in the KO cerebellum suggests a direct role of this epigenetic modification in cerebellar development.

Conditional deletion of *Mll1* [20] or *Dpy30* by the same *hGFAP-Cre* transgene expression results in partially similar phenotypes yet with important differences. The physiological phenotypes following loss of Mll1 or Dpy30 are highly similar, and include ataxia, postnatal growth retardation and death around P25-P30 (for Mll1) or P20-P27 (for Dpy30). The effects on the brain ultrastructure are also similar with hypocellular DG and expanded SVZ. However, important differences are revealed at cellular and molecular levels. While Mll1 is important for neurogenesis, but not for gliogenesis, Dpy30 is important for NPC differentiation to both lineages in a cell-intrinsic manner. Moreover, NSC self-renewal was not examined for *Mll1* KO, but the unaffected gliogenesis *in vivo* and *in vitro* suggest that NSC maintenance is probably not affected by Mll1 loss [20]. Dpy30, however, appears to be required for maintaining a functional pool and the self-renewal capacity of NSCs both *in vivo* and *in vitro*. At the molecular level, Mll1 loss leads to reduced expression *Dlx2* and *Mash1*, and this effect is not accompanied by alteration of local H3K4 methylation. Dpy30 loss, however, profoundly impaired the induction potential of several crucial genes involved in neurogenesis, accompanied by dramatic loss of global H3K4 methylation and modest reduction of H3K4me3 at affected genes. We did not detect a reduction in H3K4me3 at a tested site near the transcription start site of *GFAP* following a significant reduction in DPY30 binding at the same site upon DPY30 KD in the ReNcell VM human NPCs (Figures 4G and S4E). Based on the dependence of H3K4me3 on Dpy30 at the biochemical and genomic levels [21], we surmise that a reduction of H3K4me3 could be at a different site that would be shown by sequencing but not a single-point qPCR assay. Alternatively, we cannot rule out the possibility that other activities of DPY30 may be responsible for its regulation of GFAP expression.

These results highlight a complex relationship between the enzymatic and non-enzymatic activities of the subunits in the histone modification writer complexes. The important roles of Mll1 in regulating hematopoietic stem cells, leukemogenesis, and neurogenesis do not require its H3K4 methylation activity, and are most likely mediated by its interaction with other proteins [20, 64]. Dpy30, being a core and common subunit of all of the six Set1/Mll complexes, is indispensable for efficient H3K4 methylation throughout the genome. Our *Dpy30* KO thus allowed us to effectively perturb H3K4 methylation in this study to reveal a profound defect in NSCs deficient of this modification. Moreover, depletion of ASH2L, another common core subunit of all SET1/MLL complexes, also affected the expression of neurogenic and gliogenic genes upon treatment of differentiation conditions in the NPCs. Therefore, Dpy30 regulates NSC function most likely, or at least partially, through interacting with Ash2l and regulating the H3K4 methylation activity of one or more of the Set1/Mll subunits. In this regard, it is interesting to see the different expression pattern of the individual core and catalytic subunit of the Set1/Mll complexes at a single cell level (Figure S1B). Being the only core subunit that is preferentially expressed at NSPCs compared to their progeny, Dpy30 may hold a special position in regulating NSPC activities. The high expression of Mll3 and Mll4 in NSPCs suggests a potentially important role for the Mll3 and Mll4-mediated enhancer H3K4me1 activity, which is also regulated by the core subunits including Dpy30, in regulating NSPC activities. In addition to the Set1/Mll complex components as the major associated proteins, Dpy30 also associates with a few other proteins including the NURF chromatin-remodeling complexes and AKAP95 [14, 65]. We thus cannot exclude a role of Dpy30 in NSC regulation through interacting with these factors.

## CONCLUSION

In summary, by using both genetic inactivation in mouse and in vitro assays, our studies uncover a profound requirement of Dpy30 and its associated H3K4 methylation in sustaining the functionality of NSCs through regulating both the self-renewal and lineage differentiation capacities. This knowledge may have implications in neurodevelopmental disorders associated with mutations in the writers, erasers, and readers of H3K4 methylation.

## ACKNOWLEDGEMENTS

We thank Zhenhua Yang for technical assistance with mice dissection. This work was supported by Start-up funds from the University of Alabama at Birmingham, the University of Virginia, and NIH grant R01DK105531. HJ is a recipient of American Society of Hematology Scholar Award, American Cancer Society Research Scholar Grant, and The Leukemia & Lymphoma Society Scholar Award.

## Author Contributions

Kushani Shah: Design, Collection and/or assembly of data, Data analysis and interpretation, Manuscript writing

Gwendalyn King: Design, Data analysis and interpretation, Manuscript writing

Hao Jiang: Conception and design, Financial support, Data analysis and interpretation, Manuscript writing

## SUPPLEMENTAL FIGURES

**Supplemental Figure legends are at the end of each supplemental figures**

**Supplemental Figure 1.**
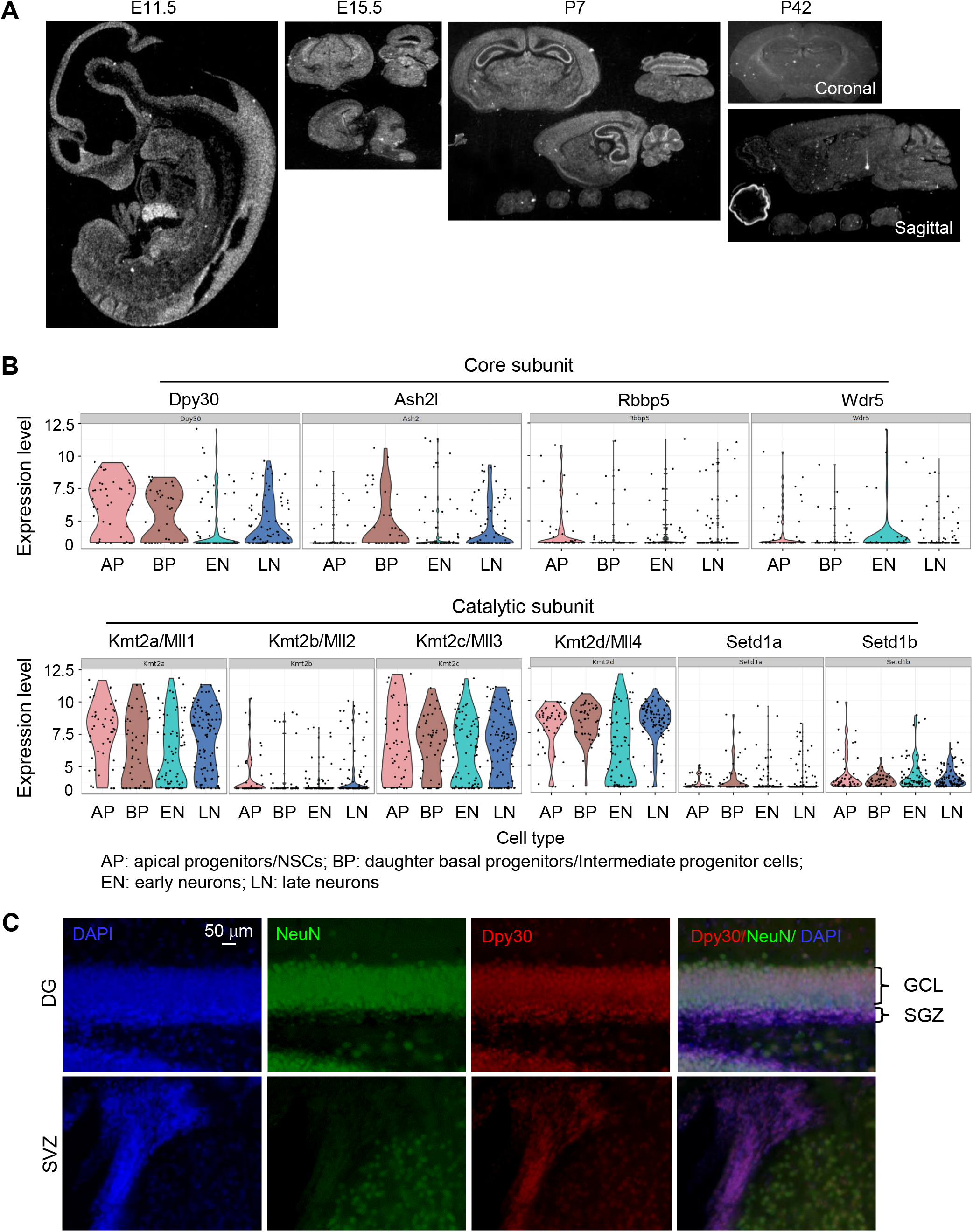
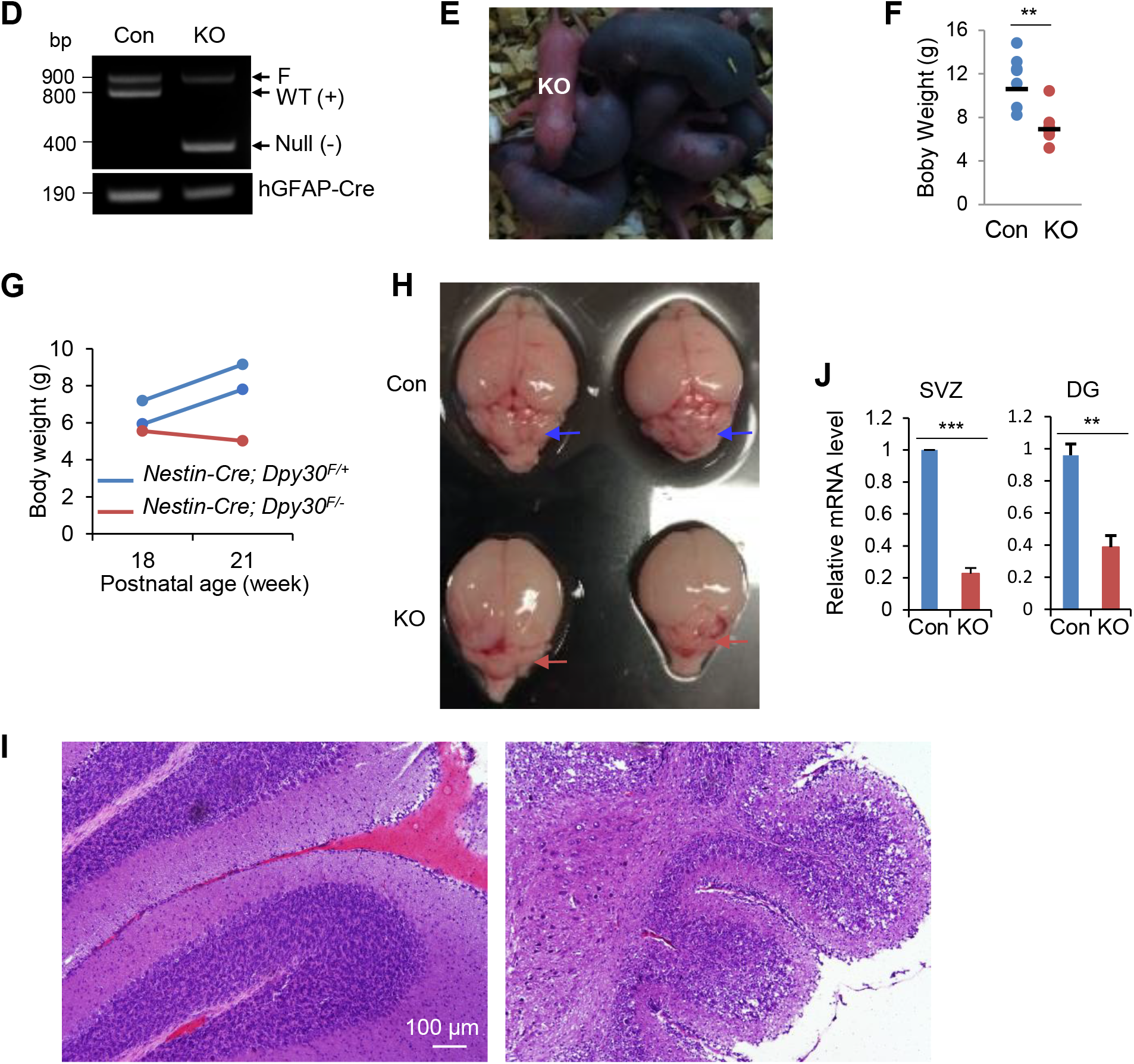
Dpy30 deficiency disrupts postnatal brain development. **(A)** Selected *in situ* hybridization images from http://www.gensat.org/bgem_ish.jsp?probe_id=3224 showing Dpy30 expression in the nervous system during development. **(B**) Violin plots showing expression level of core (top) and catalytic (bottom) subunits of the Set1/Mll complexes in indicated cell types, generated by http://genebrowser.unige.ch/science2016/ and based on single-cell RNA-seq dataset of mouse neocortical neurogenesis (Telley et al., 2016). **(C)** Co-staining of Dpy30 and NeuN in dentate (top) and SVZ (bottom) from a P21 wild type C57BL/6 mouse. Note that while Dpy30 is present in cells in SVZ, and the GCL and SGZ of DG, NeuN is only present in GCL of DG. SVZ, Subventricular Zone; SGZ, Subgranular Zone; DG, Dentate Gyrus. GCL: granular cell layer. **(D)** Genotypes of control (Con) and KO mice using genomic DNA from tail tips. **(E)** Five day old pups from the same litter photographed. Only the pink pup is KO. **(F)** Body weights of control and KO mice (n = 6 each). The short horizontal lines represent the mean. **(G)** Body weights of *Nestin-Cre; Dpy30F/+* and *Nestin-Cre; Dpy30F/-* mice are plotted at different ages. Each line represents an individual animal. **(H)** Brain dissected from P12 control and KO mice. Arrows point to the cerebellum folds. **(I)** Hematoxylin and eosin staining for the coronal cerebellum sections of P15 control and KO mice. **(J)** *Dpy30* mRNA levels in the neurogenic regions of P12 mice were determined by qPCR and normalized to *Actb*. n = 11 for SVZ, n = 3 for DG, for each genotype. Data represent the mean ± SD for (J). \*\**P* < 0.01 and \*\*\**P* < 0.001, by two-tailed Student’s t test for (E) and (J).

**Supplemental Figure 2.**
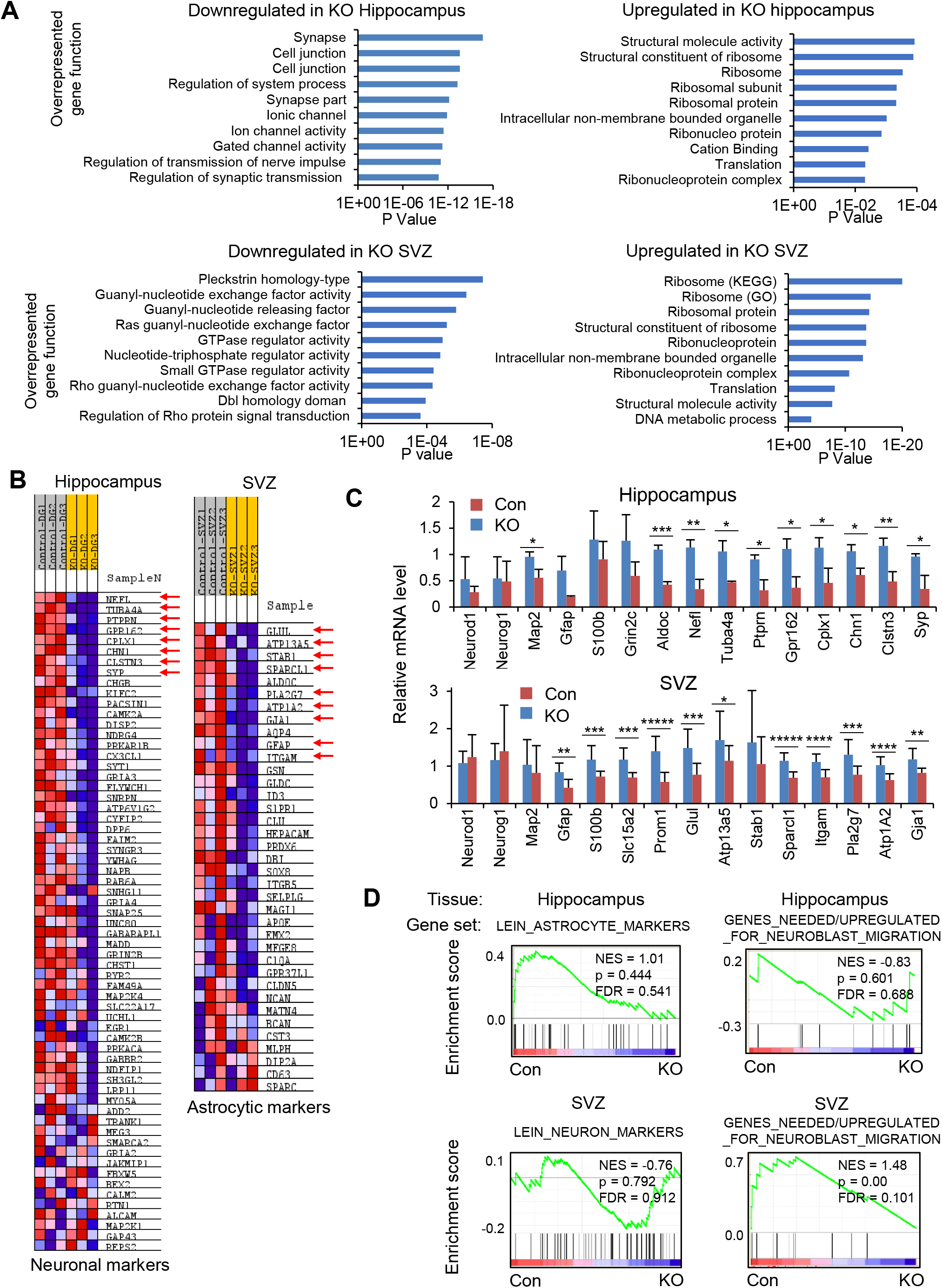
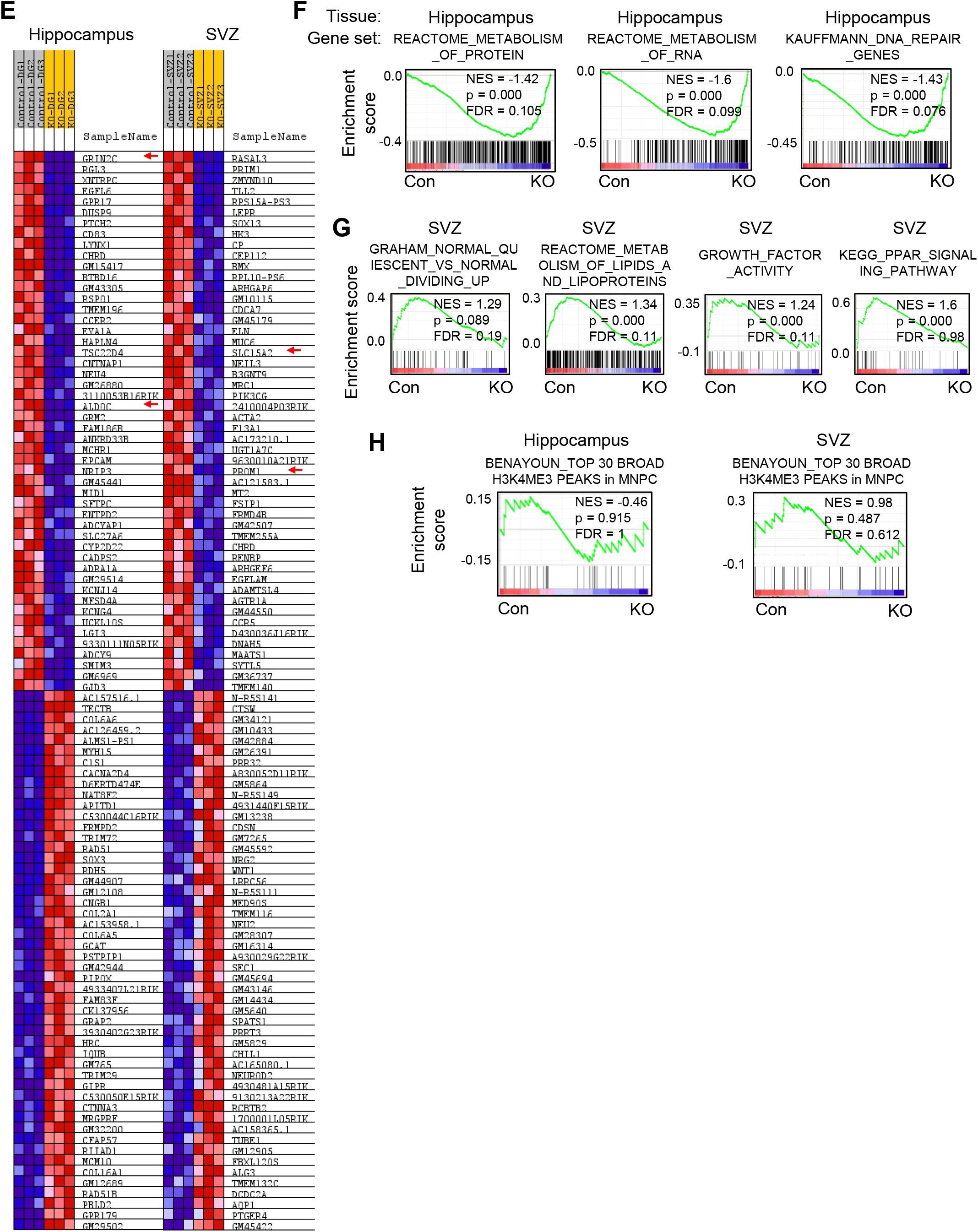
Transcriptome analyses suggest a requirement of Dpy30 for NSPC fate determination. **(A)** Top 10 GO terms enriched in genes up- and down-regulated (over 2 fold) in KO Hippocampus (top) and SVZ (bottom). n = 3 mice of each genotype for both Hippocampus and SVZ. **(B)** Heatmaps generated by GSEA showing expression changes for neuronal and astrocyte markers for Figure 2A. Genes marked with red arrows were used for analysis in (C). **(C)** Relative mRNA levels of selected genes were determined by qPCR and normalized to *Gapdh*. n = 3 mice each genetype for hippocampus. n = 12 (control) or 11 (KO) mice for SVZ. Genes include those from ‘Astrocytic marker’ and ‘Neuronal marker’ gene sets in (B) and from the top 50 up- or down-regulated genes shown in (E). **(E)** Heatmaps of the top 50 down- and upregulated genes (*P* < 0.001) following *Dpy30* KO in hippocampus (left) and SVZ (right) tissues. Genes marked with red arrows were used in (C). **(D and F-H)** GSEA on RNA-seq results obtained from SVZ and hippocampus of P12 control and KO mice. ‘Astrocyte marker’ gene set and ‘neuronal marker’ gene set (Lein et al., 2007) and curated neuroblast migration gene set were used for (D). ‘Active NSC’ gene sets and ‘quiescent NSC’ gene sets (Codega et al., 2014) were used for (F) and (G), respectively. H3K4me3 broad peaks in mNPC gene set (Benayoun et al., 2014) were used for (H). Data represent the mean ± SD in (C). \**P <* 0.05, \*\**P* < 0.01, \*\*\**P* < 0.001, \*\*\*\**P* < 0.0001, and \*\*\*\*\**P* < 0.00001, by two-tailed Student’s t test for (C).

**Supplemental Figure 3.**
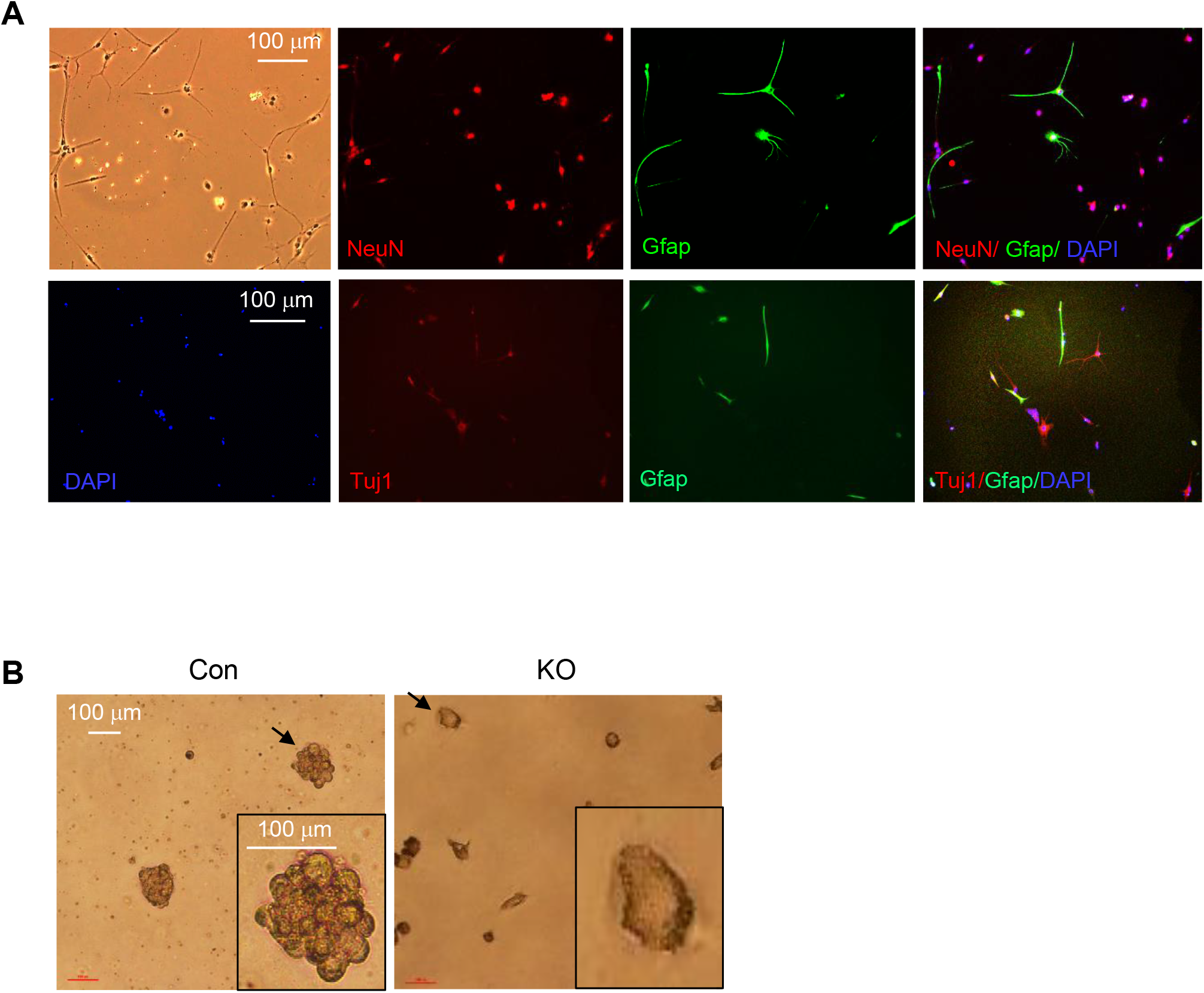
Dpy30 is required for sustaining the self-renewal and proliferation of NSPCs. **(A)** Images of differentiated neurospheres obtained from SVZ of control mice. NeuN (top) and Tuj1 (bottom) for neurons. Gfap for glia. Representatives of 3 control mice each for NeuN and Gfap staining, and 3 control mice for Tuj1 and Gfap staining. **(B)** Neurospheres in culture formed by cells extracted from DG of control mouse (left) and lack of neurospheres in culture from KO (right). Imaged at day 14 in proliferation media. Inset shows magnified image of the cells indicated by black arrows. Representatives of 2 mice for each genotype.

**Supplemental Figure 4.**
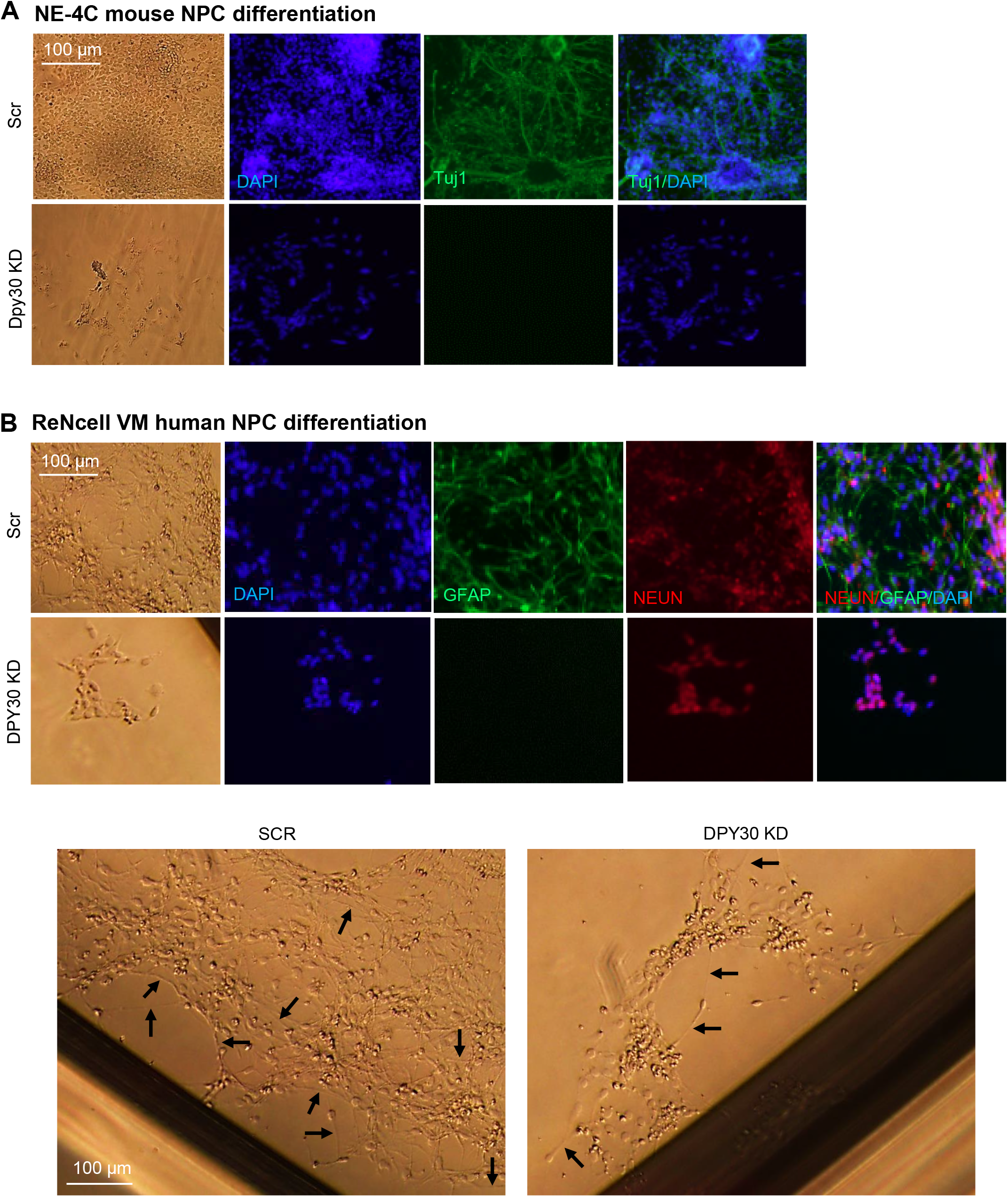
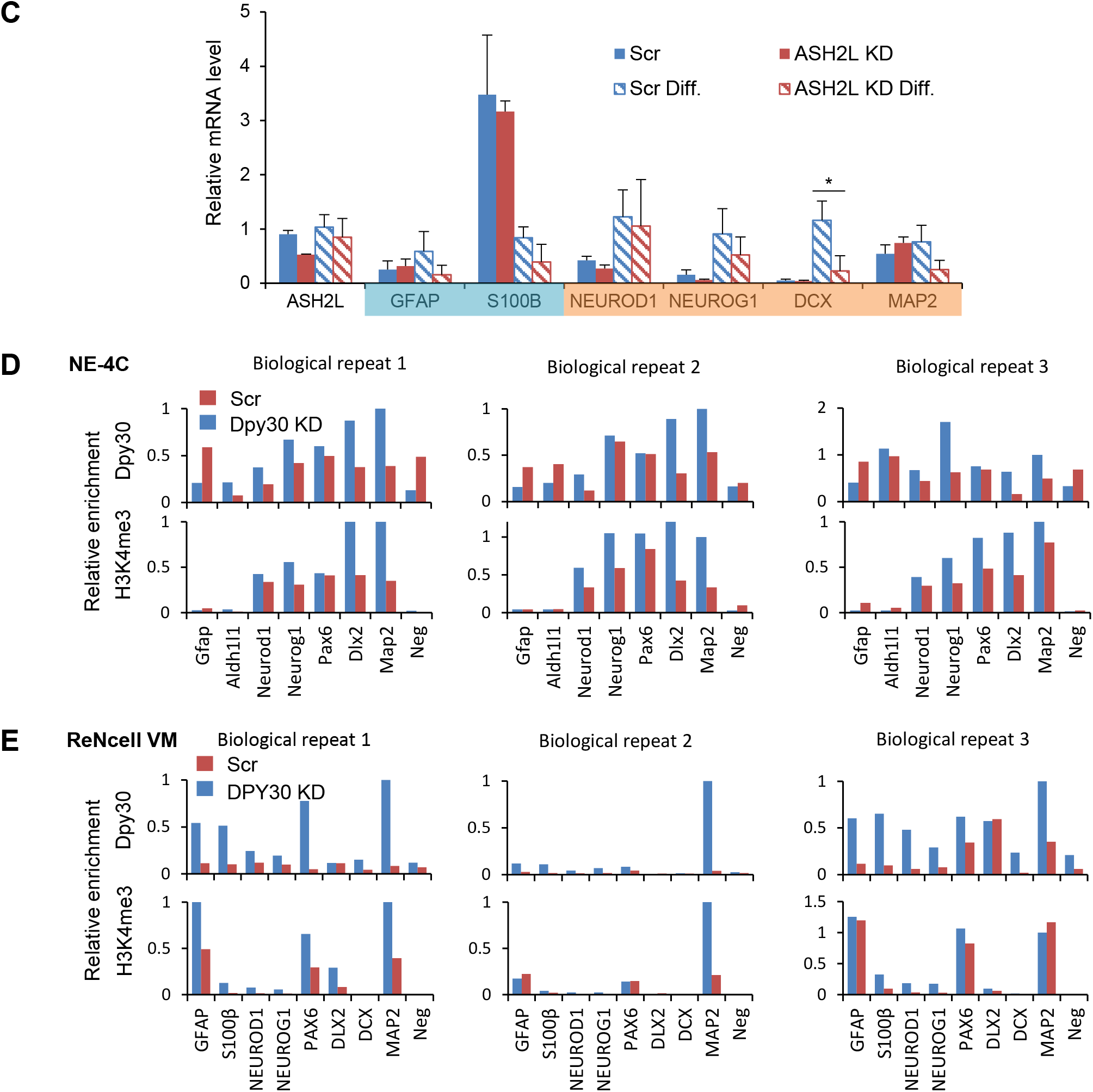
Core subunits of the Set1/Mll complexes regulate NPC differentiation and lineage gene induction. **(A)** Differentiation of NE-4C mouse NPCs. Staining performed at Day 10 in differentiation media. Tubulin β-III (Tuj1) stains neurons. Representative of 3 biological repeats. **(B)** Differentiation of ReNcell VM human NPCs. Staining performed at Day 10 in differentiation media. Top, NEUN stains neurons and GFAP stains astrocytes. Bottom, bright field image showing long neuronal fibers (arrows). Representative of 3 biological repeats. **(C)** Relative mRNA levels were determined and normalized to *GAPDH* for indicated genes upon ASH2L KD with and without treatment of differentiation conditions in ReNcell VM cells. One out of the 3 biological repeat ‘Scr Differentiated’ was set to 1 for all genes. The shade over the gene names represents groups involved in different processes: aqua, gliogenesis; orange, neurogenesis. **(D-E)** Individual biological repeats shown for Figure 4C and 4G in (D) and (E), respectively. Data represent the mean ± SD from 3 biological repeats for (C). \**P* < 0.05, by two factor ANOVA analysis followed with post hoc *t*-Test for (C).

## SUPPLEMENTAL TABLES

**Supplemental Table 1. Normalized transcript counts in hippocampus and SVZ.**

There are 2 tabs in this file, one tab for SVZ and the other for hippocampus. Listed are normalized transcript counts by RNA-seq for all genes in individual samples from 3 control mice and 3 KO mice.

**Supplemental Table 2. Significantly changed genes in hippocampus and SVZ.**

There are 4 tabs in this file, which, respectively, list genes that were: significantly downregulated over 2 fold in the KO hippocampus (“Hippo DN in KO>2 fold p<0.05” tab), significantly upregulated over 2 fold in the KO (“Hippo UP in KO>2 fold p<0.05” tab), significantly downregulated over 2 fold in the KO SVZ (“SVZ DN in KO>2 fold p<0.05” tab), significantly upregulated over 2 fold in the KO SVZ (“SVZ UP in KO>2 fold p<0.05” tab), all compared to the corresponding control tissue. Also shown are their mean expression levels (normalized reads) among the 3 different samples of each genotype, the p-value and FDR, as determined by DE-seq analyses of the RNA-seq results. The genes were sorted by fold change.

**Supplemental Table 3.**
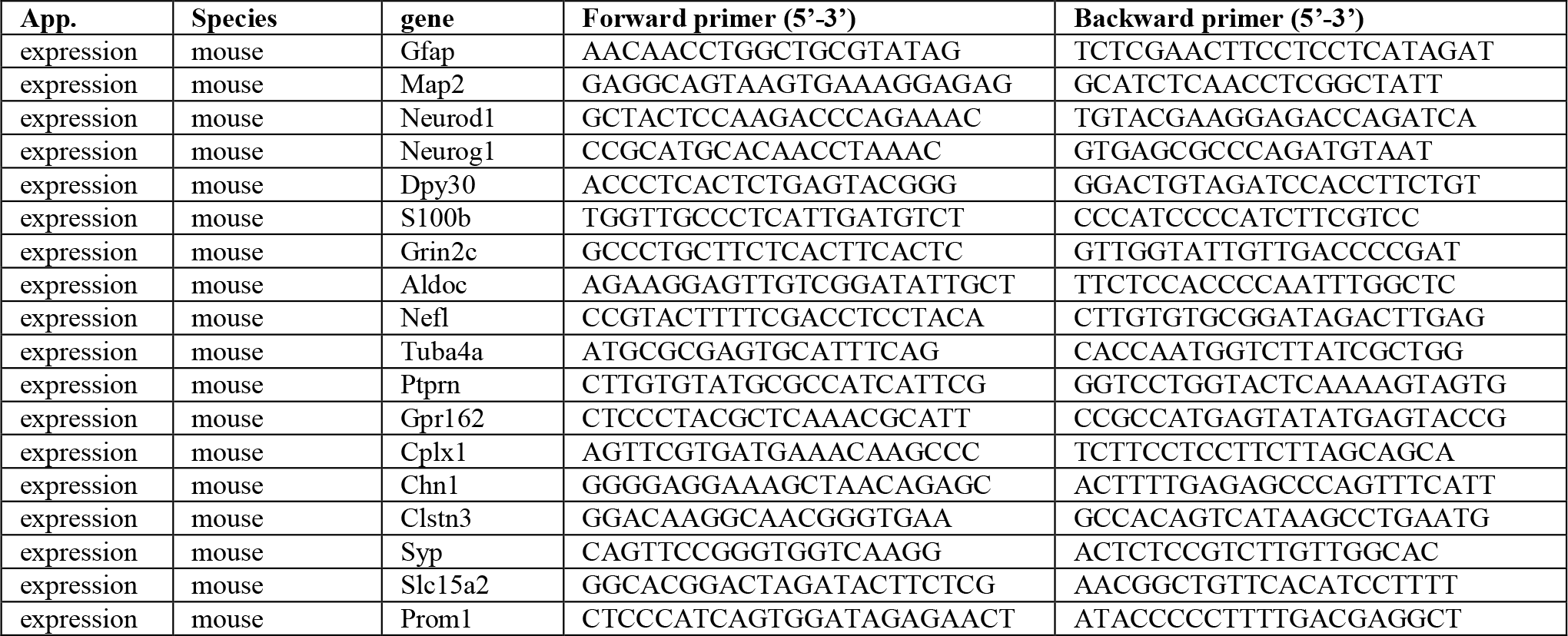
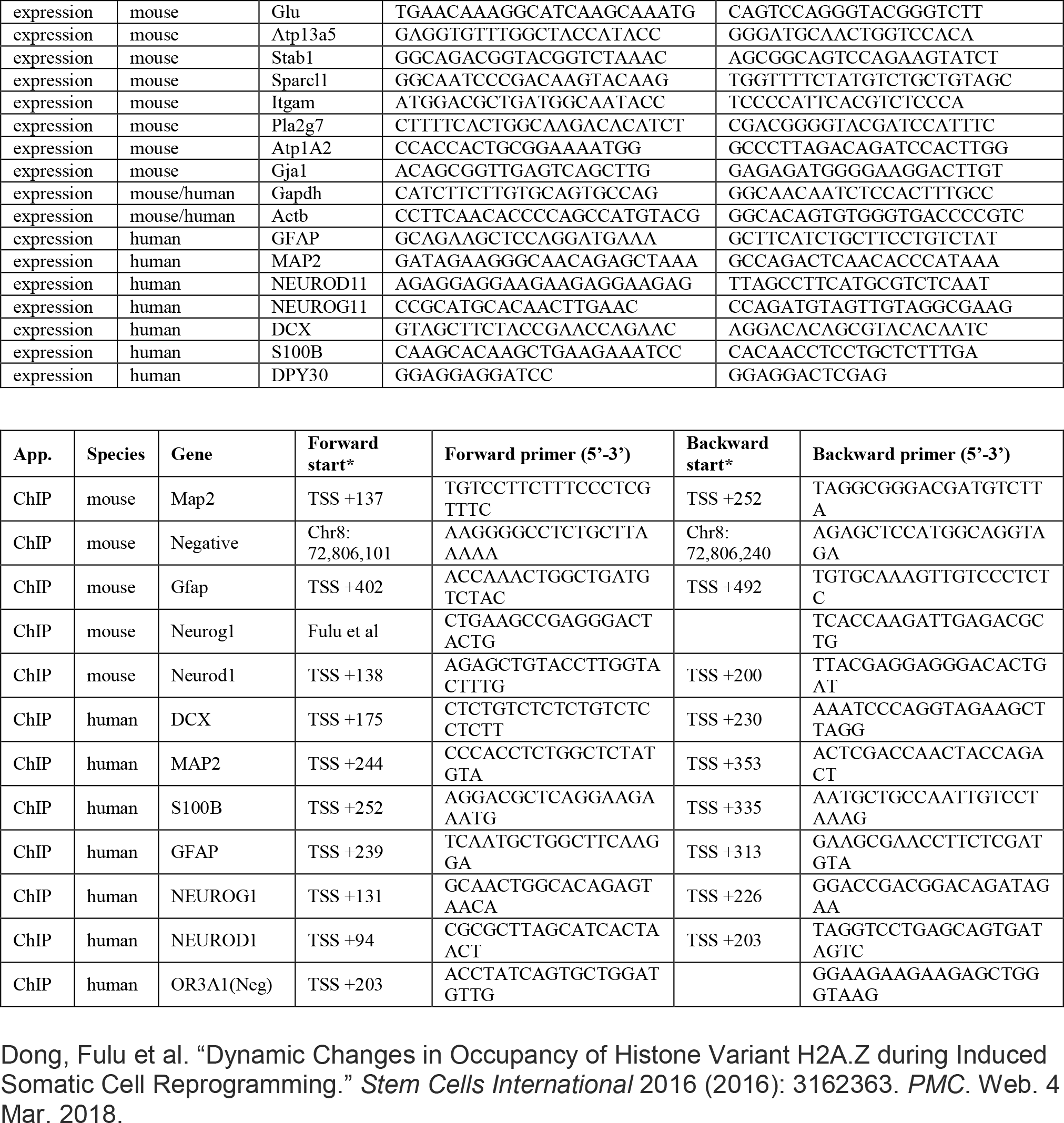
qPCR Primers. Dong, Fulu et al. “Dynamic Changes in Occupancy of Histone Variant H2A.Z during Induced Somatic Cell Reprogramming.” *Stem Cells International* 2016 (2016): 3162363. *PMC*. Web. 4 Mar. 2018.

## SUPPLEMENTAL VIDEO

**Supplemental video 1. Ataxia-like symptoms of the *Dpy30* KO mice.** Two mice on the left are *hGFAP-Cre; Dpy30*^*F/-*^ littermates with different severity. On the right is the *hGFAP-Cre; Dpy30*^*F/+*^ control littermate for comparison.

Author Contributions
Kushani Shah: Design, Collection and/or assembly of data, Data analysis and interpretation, Manuscript writing Gwendalyn King: Design, Data analysis and interpretation, Manuscript writing Hao Jiang: Conception and design, Financial support, Data analysis and interpretation, Manuscript writing

